# Mitochondrial dysfunction is signaled to the integrated stress response by OMA1, DELE1 and HRI

**DOI:** 10.1101/715896

**Authors:** Xiaoyan Guo, Giovanni Aviles, Yi Liu, Ruilin Tian, Bret A. Unger, Yu-Hsiu T. Lin, Arun P. Wiita, Ke Xu, M. Almira Correia, Martin Kampmann

**Affiliations:** Institute for Neurodegenerative Disease, University of California, San Francisco, San Francisco, CA, USA; Chan Zuckerberg Biohub, San Francisco, CA, USA; Department of Cellular and Molecular Pharmacology, University of California, San Francisco, San Francisco, CA, USA; Biophysics Graduate Program, University of California, San Francisco, San Francisco, CA, USA; Department of Chemistry, University of California, Berkeley, Berkeley, CA, USA; Department of Laboratory Medicine, University of California, San Francisco, San Francisco, CA, USA; Department of Pharmaceutical Chemistry, University of California, San Francisco, San Francisco, CA, USA; Department of Bioengineering and Therapeutic Sciences, University of California, San Francisco, San Francisco, CA, USA; The Liver Center, University of California, San Francisco, San Francisco, CA, USA; Department of Biochemistry and Biophysics, University of California, San Francisco, San Francisco, CA, USA

## Abstract

In mammalian cells, mitochondrial dysfunction triggers the integrated stress response (ISR), in which eIF2α phosphorylation upregulates the transcription factor ATF4. However, how mitochondrial stress is relayed to the ISR is unknown. We found that HRI is the eIF2α kinase necessary and sufficient for this relay. Using an unbiased CRISPRi screen, we identified factors upstream of HRI: OMA1, a mitochondrial stress-activated protease, and DELE1, a little-characterized protein we found to be associated with the inner mitochondrial membrane. Mitochondrial stress stimulates the OMA1-dependent cleavage of DELE1, leading to its accumulation in the cytosol, where it interacts with HRI and activates its eIF2α kinase activity. Blockade of the OMA1-DELE1-HRI pathway is beneficial during some, but not all types of mitochondrial stress, and leads to an alternative response that induces specific molecular chaperones. Therefore, this pathway is a potential therapeutic target enabling fine-tuning of the ISR for beneficial outcomes in diseases involving mitochondrial dysfunction.

## Main Text

Mitochondria are essential organelles in eukaryotic cells that play a central role in energy homeostasis, metabolism and signaling. Mitochondrial function is challenged by disease conditions, environmental toxins and aging^1-3^. Mitochondrial dysfunction elicits cellular stress responses. The precise molecular mechanisms underlying some mitochondrial stress responses in model organisms have been elucidated, particularly for the mitochondrial unfolded protein response (mito-UPR) in *C. elegans*, which is mediated by the transcription factor ATFS-1^4^. In mammalian cells, ATF5 is proposed to play an equivalent role^5^. Mammalian cells also have additional pathways that mediate a mito-UPR in response to disruption of mitochondrial protein homeostasis^6-10^.

However, the overwhelming signature in the response of human cells to disruption of a broad range of mitochondrial functions is activation of the integrated stress response (ISR), as previously published^11-13^ and confirmed by us (Extended Data Fig. 1). The ISR is mediated through phosphorylation of the translation initiation factor eIF2α under various stress conditions, which are sensed by four eIF2α kinases, PKR, GCN2, PERK and HRI^14-17^. Phosphorylation of eIF2α reduces global protein translation but selectively increases translation of certain poorly translated mRNAs with upstream open reading frames (uORFs), including the mRNA encoding ATF4, the master transcriptional regulator of the ISR. Currently, it is unknown how mitochondrial dysfunction triggers the ISR. Here, we uncover the underlying molecular mechanism in human cells: mitochondrial stress leads to cleavage and cytosolic accumulation of the little-characterized mitochondrial protein DELE1, which then interacts with HRI and activates it to trigger the ISR. This response can be beneficial for some types of mitochondrial pertubation, but maladaptive for others, and its blockade induces an alternative transcriptional program.

### The eIF2α kinase HRI signals mitochondrial stress to the Integrated Stress Response

To characterize how mitochondrial dysfunction is signalled to the ISR, we constructed a translational reporter for the ISR, in which uORF1 and 2 of ATF4 precede the coding sequence of the mApple fluorescent protein (Fig. 1a). To control for transcription of this reporter from the CMV promoter, we generated a second reporter in which CMV directs the transcription of EGFP, without uORFs. We introduced these reporters into human HEK293T cells, along with the machinery necessary for CRISPR interference (CRISPRi), which utilizes catalytically dead Cas9 fused to a transcriptional repressor domain (KRAB) to knock down expression of endogenous genes as directed by a single guide RNA (sgRNA)^18,19^ (Fig. 1a).

**Figure 1.**
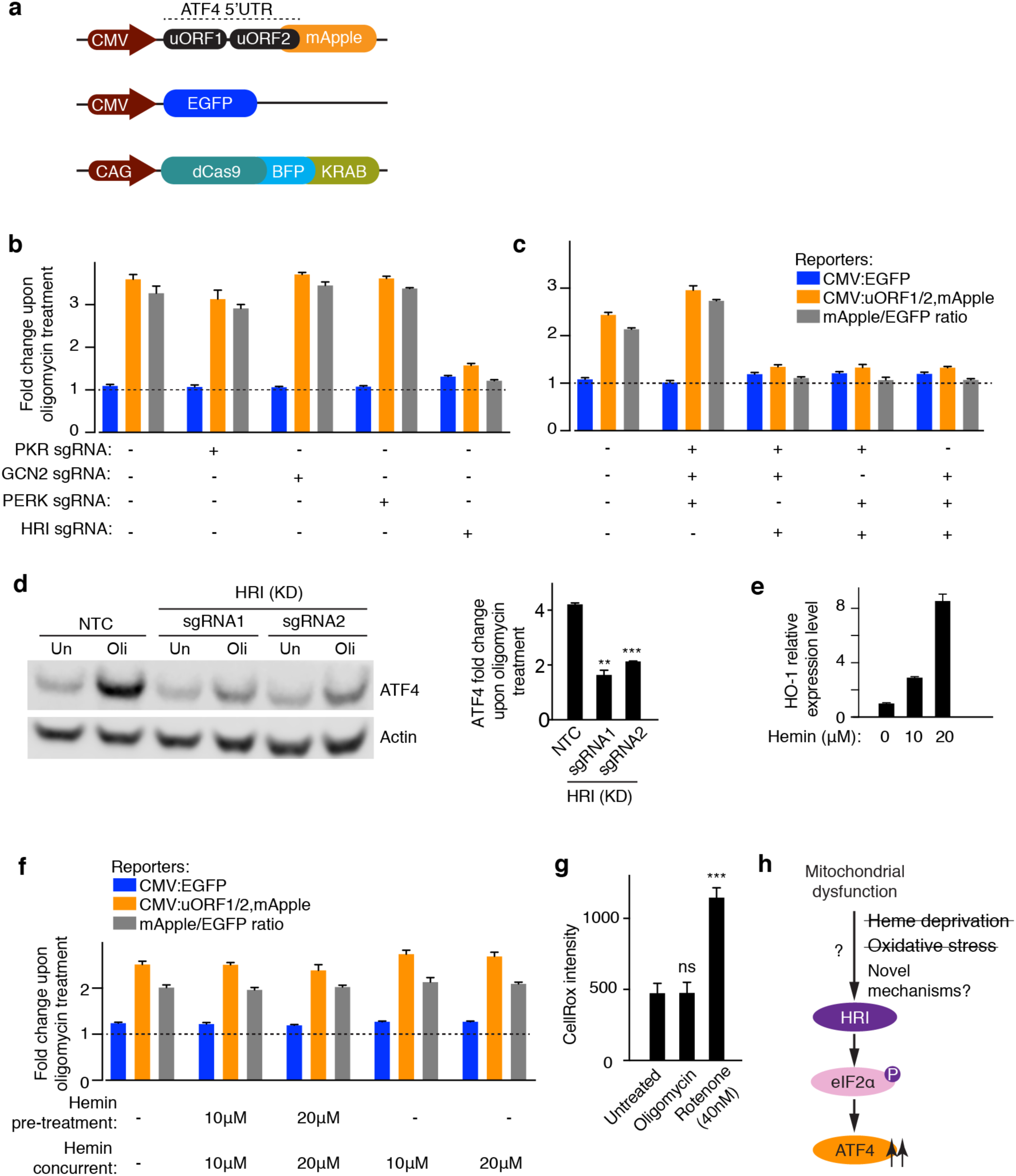
The eIF2α kinase HRI relays mitochondrial stress to the integrated stress response. **(a)** Reporters and CRISPRi constructs. The ATF4 translational reporter includes the upstream open reading frames (uORFs) 1 and 2 of the ATF4 5’ untranslated region (5’UTR) followed by mApple replacing the ATF4 coding sequence. The transcription of the ATF4 translational reporter is under the control of the constitutive CMV promoter. A secondary reporter with EGFP directly driven by CMV serves as a transcriptional control. Catalytically dead Cas9 (dCas9) fused to BFP and a transcriptional repressor domain (KRAB) is under the control of the CAG promoter to knock down gene expression. These constructs were introduced into human HEK293T cells to generate the reporter cell line. **(b**,**c)** HRI is necessary and sufficient to activate ATF4 translation upon mitochondrial stress. Reporter cells expressing either single sgRNAs (b) or triple sgRNAs (c) targeting the indicated eIF2α kinases were exposed to 1.25 ng/mL oligomycin for 16 h before measuring reporter levels by flow cytometry. The reporter fold change (mean ± s.d., *n* = 3 culture wells) is the ratio of median fluorescence values for oligomycin over untreated samples. **(d)** Western blot of endogenous ATF4. Cells expressing a non-targeting control sgRNA (NTC) or sgRNAs targeting HRI were untreated or treated with 1.25 ng/mL of oligomycin for 16 h. *Left*, representative blot; *Right*, quantification of *n* = 2 Western blots (mean ± s.d., \*\**P* < 0.01,\*\*\**P* < 0.001, unpaired *t* test). **(e)** Expression of the heme-induced gene HO-1 in HEK293T cells incubated for 24 h with the indicated concentrations of hemin measured by quantitative RT-PCR (mean ± s.d., *n* = 3 technical replicates). **(f)** Heme supplementation does not abolish ATF4 induction, indicating that HRI is not activated via heme depletion during mitochondrial stress. Reporter cells were untreated or treated with the indicated concentrations of hemin for 24 h before a 16 h treatment with 1.25 ng/mL oligomycin in the presence or absence of hemin, and reporter levels (mean ± s.d., *n* = 3 culture wells) were quantified as in (b,c). **(g)** Oligomycin treatment used in this study does not induce reactive oxygen species (ROS). HEK293T cells were treated with 1.25 ng/mL oligomycin or 40 nM rotenone for 16 h and ROS levels were quantified by flow cytometry using the CellROX reagent (mean ± s.d., *n* = 3 culture wells, \*\*\**P* < 0.001, ns, not significant, two-tailed unpaired *t* test). **(h)** A model summarizing the findings described in this Figure.

We validated this ISR reporter cell line using thapsigargin to induce endoplasmic reticulum (ER) stress. Thapsigargin activated the reporter, and as expected, this induction was blocked by knockdown of PERK (Extended Data Fig. 2a,b), the eIF2α kinase known to signal ER stress to the ISR. The reporter was also activated by a broad range of mitochondrial stresses, including pharmacological inhibitors of different mitochondrial functions (the mitochondrial ribosome inhibitor doxycycline, the electron transport chain inhibitors antimycin A and rotenone, and the ATP synthase inhibitor oligomycin, Extended Data Fig. 2c) and genetic knockdown of proteins required for mitochondrial protein homeostasis (HSPD1 and LONP1) and mitochondrial ribosomal proteins (MRPL17 and MRPL22) (Extended Data Fig. 2d). We opted to use oligomycin as the primary mitochondrial stressor in this study, since it substantially induced the ATF4 translational reporter, but not transcription from the CMV promoter (Extended Data Fig. 2c).

Next, we asked whether mitochondrial stress triggers the ISR via one of the known eIF2α kinases. Intriguingly, knockdown of HRI, but none of the other three eIF2α kinases, significantly reduced the oligomycin-induced activation of the ISR reporter (Fig. 1b). We also knocked down all triple combinations of eIF2α kinases simultaneously and showed that HRI is sufficient to mediate oligomycin induction of the ISR reporter (Fig. 1c). Western blot of endogenous ATF4 further validated that HRI is responsible for ATF4 up-regulation in response to mitochondrial stress (Fig. 1d).

This finding raised the question of how dysfunction in the mitochondria is sensed by the cytosolic factor HRI. Since the canonical mechanism of HRI activation is a decrease of heme levels^20^ and since key steps in heme biosynthesis occur in the mitochondria^21^, we hypothesized that heme depletion could be the signal by which HRI is activated upon mitochondrial dysfunction. We reasoned that if mitochondrial stress activated HRI through heme deprivation, hemin supplementation would block the oligomycin-induced ATF4 up-regulation. We confirmed that hemin could enter the reporter cells, as evidenced by the induction of the heme-inducible gene HO-1 (Fig. 1e). However, hemin supplementation neither before nor during oligomycin treatment reduced ISR induction, indicating that heme depletion is not the signal transmitting mitochondrial stress to HRI (Fig. 1f).

Our next hypothesis was that reactive oxygen species (ROS), which can be produced by dysfunctional mitochondria, may act as the signal by oxidizing and thereby activating HRI, via another previously described mechanism^22,23^. However, we found that, unlike rotenone treatment, the oligomycin concentration sufficient to trigger the ISR did not lead to a detectable ROS elevation (Fig. 1g), ruling out ROS as the relevant signal under these conditions. These findings suggest a novel mechanism of HRI activation upon mitochondrial stress (Fig. 1h).

### A CRISPRi screen identifies the mitochondrial protein DELE1, which acts upstream of HRI

To identify the molecular players mediating ISR activation upon mitochondrial stress, we conducted a large-scale CRISPRi screen. ISR reporter cells were infected with pooled sgRNA libraries targeting 7,710 protein-coding genes (including signaling proteins, protein homeostasis factors, and mitochondrial proteins; the libraries were a subset selected from our genome-wide next-generation CRISPRi libraries^24^). Cells were treated with oligomycin or left untreated for 16 hours, and cell populations were sorted based on reporter activity (the ratio between mApple and EGFP) using fluorescence-activated cell sorting (Fig. 2a). The frequencies of cells expressing each sgRNA in the different populations were determined by targeted next-generation sequencing, and genes whose knockdown significantly altered reporter activity were detected using our previously described quantitative framework^25,26^. Phenotypes for all targeted genes are provided in Supplemental Table 3. We focused on the category of hit genes whose knockdown reduced reporter activity in the oligomycin-treated but not untreated cells. HRI was the second strongest hit in this category (Fig. 2b), consistent with the results from our previous experiments (Fig. 1bc). The strongest hit in this category was the little-characterized gene DELE1 (KIAA0141) (Fig. 2b). Using individually cloned sgRNAs targeting DELE1, we validated that DELE1 knockdown inhibited the induction of the ISR reporter and endogenous ATF4 in response to oligomycin, with stronger knockdown resulting in greater inhibition (Fig. 2cde).DELE1 was not required for ISR activation in response to ER stress (Extended Data Fig. 3), suggesting that it acts specifically in signaling mitochondrial stress to the ISR.

**Fig. 2.**
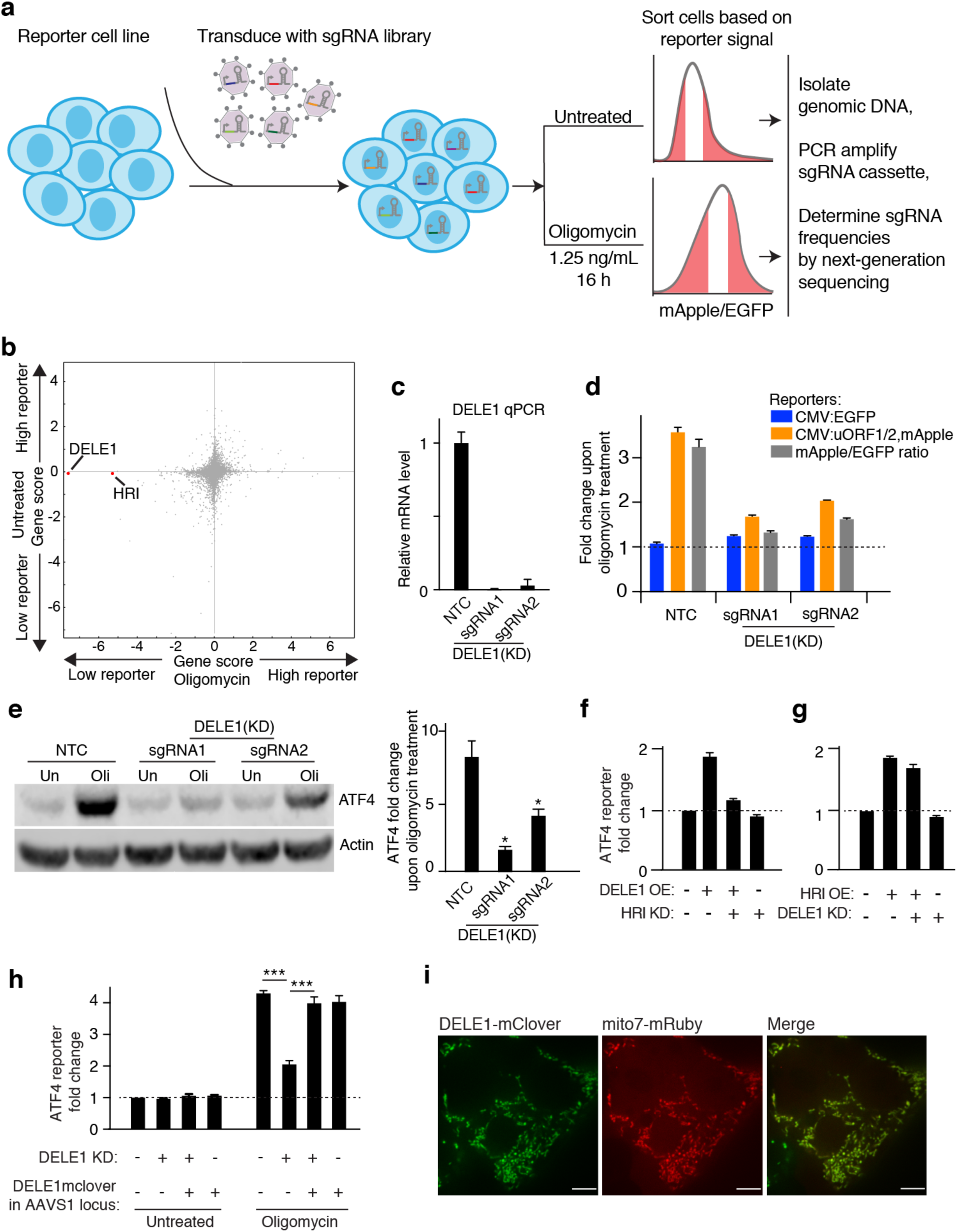
A CRISPRi screen identifies the mitochondrial protein DELE1, which acts upstream of HRI. **(a)** Strategy for the CRISPRi screen. Reporter cells transduced with an sgRNA library were cultured for 16 h untreated or treated with 1.25 ng/mL oligomycin before FACS sorting. The top and bottom 30% of cells based on reporter ratio were collected from each population for targeted next-generation sequencing. **(b)** Comparison of gene scores (defined in Methods) for the 7,710 genes targeted in the CRISPRi screen (grey dots) in untreated and oligomycin-treated conditions. Highlighted in red are two genes (HRI and DELE1) knockdown of which significantly reduces ATF4 in the oligomycin treated condition, but not in the untreated condition. **(c)** Knockdown of DELE1 by CRISPRi by two sgRNAs, quantified by qPCR (mean ± s.d., *n* = 3 technical replicates). **(d)** Reporter activation was measured as in Fig. 1b,c in cells expressing a non-targeting control sgRNA (NTC) or sgRNAs targeting DELE1 (mean ± s.d., *n* = 3 culture wells). **(e)** Western blot of endogenous ATF4. Cells expressing a non-targeting control sgRNA (NTC) or sgRNAs targeting DELE1 were untreated or treated 1.25 ng/mL of oligomycin for 16 h. *Left*, representative blot; *Right*, quantification of *n* = 2 blots (mean ± s.d., \**P* < 0.05, two-tailed unpaired *t* test). **(f)** DELE1 was overexpressed (OE) using transient transfection in reporter cells expressing sgRNA to knockdown (KD) HRI, or no sgRNA, and reporter activity was quantified as in Fig. 1b,c (mean ± s.d., *n* = 3 culture wells). **(g)** HRI was overexpressed (OE) using transient transfection in reporter cells expressing sgRNA to knockdown (KD) DELE1, or no sgRNA, and reporter activity was quantified as in Fig. 1b,c (mean ± s.d., *n* = 3 culture wells). **(h)** A transgene of DELE1 C-terminally tagged with mClover stably expressed from the AAVS1 safe-harbor locus is not sufficient to induce the ATF4 reporter in the absence of oligomycin, but can rescue the DELE1 knockdown phenotype in response to oligomycin. Reporter activity was quantified as in Fig. 1b,c (mean ± s.d., *n* = e culture wells, \*\*\**P* < 0.001, two-tailed unpaired *t* test). **(i)** Co-localization of stably expressed DELE1-mClover with the mitochondrial-targeted mRuby (Mito7-mRuby). Scale bar, 7 μm.

DELE1 was previously identified as a mitochondrial protein with a role in death receptor-mediated apoptosis^27^. To further characterize DELE1, we tested four commercially available antibodies against DELE1, but none detected endogenous DELE1 (Extended Data Fig. 4). Therefore, we tagged DELE1 C-terminally with mClover to enable its further characterization. We found that transient transfection with a DELE1-mClover plasmid led to a substantial overexpression compared to endogenous DELE1 (Extended Data Fig. 5) and was sufficient to induce the ATF4 reporter, and this induction required HRI (Fig. 2f). Similarly, overexpression of HRI also induced the ATF4 reporter, but in a DELE1 independent manner, indicating that DELE1 acts upstream of HRI in this pathway (Fig. 2f,g).

**Fig. 3.**
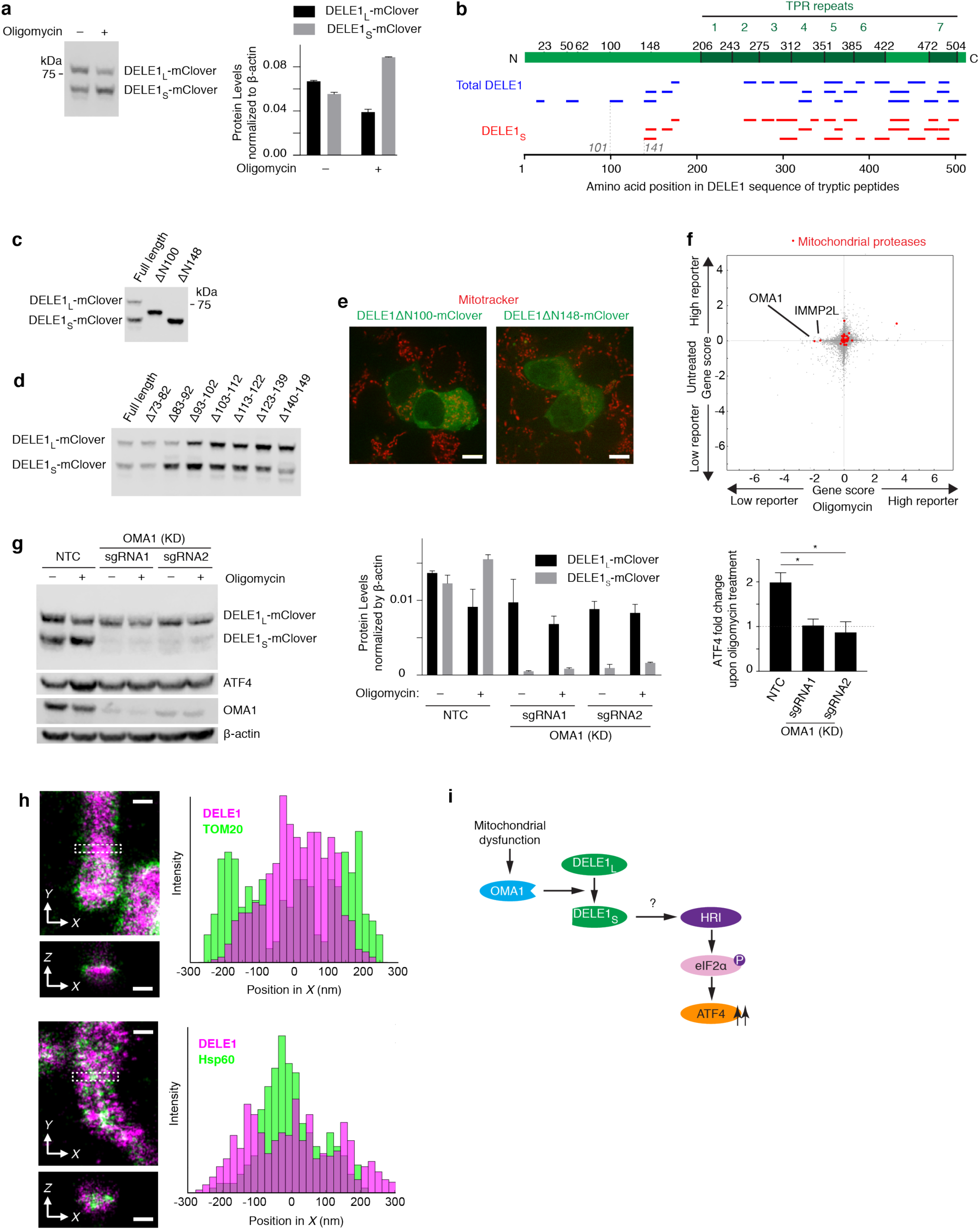
A cleaved form of DELE1 accumulates upon mitochondrial stress in an OMA1-dependent manner. **(a)** A short form of DELE1 (DELE1_S_) accumulates upon mitochondrial stress. Cells stably expressing DELE1-mClover were treated with 1.25 ng/mL of oligomycin for 16 h or left untreated, and DELE1-mClover was detected using an anti-GFP antibody. *Left*, representative blot. *Right*, quantification of *n* = 2 blots (mean ± s.d.). **(b)** Tryptic peptides of total DELE1 (blue) or DELE1_S_ (red) detected by mass spectrometry mapped to the amino acid (aa) sequence of DELE1. DELE1_S_ lacks peptides derived from sequences N-terminal to aa 141. **(c)** Western blot from cells transiently expressing full-length DELE1-mClover and truncation constructs ΔN100 and ΔN148, missing the N-terminal 100 or 148 amino acids, respectively. **(d)** Western blot from cells transiently expressing full-length DELE1-mClover and truncation constructs lacking the indicated amino acids. **(e)** Lack of co-localization of transiently expressed DELE1ΔN100-mClover and DELE1ΔN148-mClover (green) with the mitochondrial stain Mitotracker (red). Scale bar, 7 μm. **(f)** Comparison of gene scores (defined in Methods) for the 7,710 genes targeted in the CRISPRi screen (grey dots) in untreated and oligomycin-treated conditions. Highlighted in red are mitochondrial proteases, of which OMA1 and IMMP2L are the top two hits knockdown of which significantly reduces ATF4 in the oligomycin treated condition, but not in the untreated condition. **(g)** *Left*, Representative Western blot of DELE1-mClover, ATF4 and OMA1 in cells expressing non-targeting control sgRNAs (NTC) or sgRNAs for OMA1 knockdown, which were treated with 1.25 ng/mL of oligomycin for 16 h or left untreated. *Right*, quantification of *n* = 2 blots (mean ± s.d.). **(h)** Two-color 3D-STORM super-resolution images of stably expressed DELE1-mClover (magenta) in combination with the outer mitochondrial membrane marker TOM20 (*top*) or the mitochondrial matrix protein Hsp60 (*bottom*). For each pair, a virtual cross-section and a spatial intensity distribution are shown for the boxed area. Scale bars: 250 nm. **(i)** A model summarizing the findings so far.

**Fig. 4.**
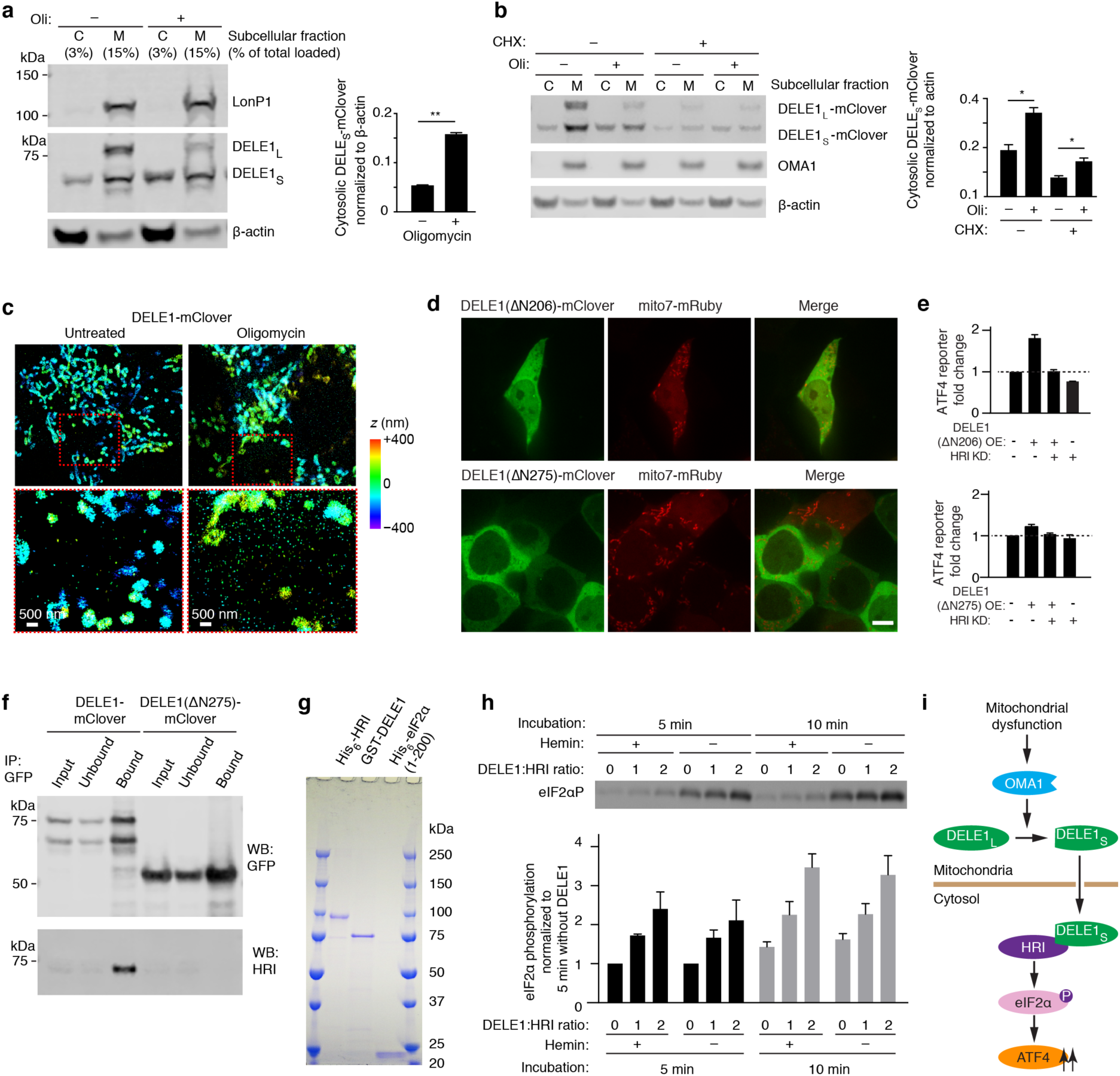
Cytosolic DELE1 physically interacts with and activates HRI. **(a)** DELE_L_ is exclusively localized in the mitochondria, while DELE1_S_ accumulates in the cytosol during mitochondrial stress. Biochemical fractionation of cells stably expressing DELE1-mClover that were either treated with 1.25 ng/mL oligomycin (Oli) for 16 h or left untreated. *Left*, representative Western blot of the cytosolic (C) and mitochondrial (M) fractions. β-actin and LonP1 were probed as markers for cytosol and mitochondria, respectively. *Bottom*, quantification of the cytosolic DELE1_S_-mClover from *n* = 2 blots (mean ± s.d., \*\**P* < 0.001, two-tailed unpaired *t* test). **(b)** Accumulation of DELE1_S_ does not depend on protein synthesis. Biochemical fractionation of cells stably expressing DELE1-mClover that were treated with 1.25 ng/mL oligomycin (Oli) for 4 h or left untreated, either in the presence or absence of 20 μg/mL cycloheximide (CHX), which inhibits cytosolic ribosomes. *Left*, representative Western blot of the cytosolic (C) and mitochondrial (M) fractions. β-actin and OMA1 were probed as markers for cytosol and mitochondria, respectively. *Right*, quantification of the cytosolic DELE1_S_-mClover from *n* = 2 blots (mean ± s.d., \*\**P* < 0.001, two-tailed unpaired *t* test). **(c)** Increased detection of DELE1-mClover outside the mitochondria upon oligomycin treatment. 3D-STORM super-resolution images of stably expressed DELE1-mClover (colors indicating depth in the z dimension) in untreated cells (*left*) and cells treated with 1.25 ng/mL oligomycin for 16 h (*right*). Areas boxed in red in the top panels are shown in higher magnification in the bottom panels. **(d)** Transiently expressed DELE1-mClover constructs lacking N-terminal 206 or 275 amino acids are localized in the cytosol. Micrographs of cells also transfected with mitochondrial-targeted mRuby (Mito7-mRuby). Scale bar, 7 μm. **(e)** Transient overexpression (OE) of the cytosolically localized DELE1(ΔN206) construct is sufficient to induce ATF4 reporter, while DELE1(ΔN275) no longer induces the ATF4 reporter. Knockdown (KD) of HRI blocks reporter activation. Reporter quantified as in Fig. 1b,c (mean ± s.d., *n* = 3 culture wells). **(f)** Co-immunoprecipitation of HRI with transiently expressed full-length DELE1-mClover but not with a construct lacking the N-terminal 275 amino acids, DELE1(ΔN275)-mClover. **(g)** Purified recombinant HRI, DELE1 and eIF2α. 800 ng of each recombinant protein was subjected to SDS-PAGE and stained with Coomassie blue. **(h)** *In vitro* eIF2α kinase assay of recombinant HRI in the presence of increasing concentrations of recombinant DELE1 and in the presence or absence of 5 µM hemin. *Top*, representative Western blot of phospho-eIF2α. Bottom, quantification (mean ± s.e.m., *n* = 3 reactions). **(i)** Model: OMA1, DELE1 and HRI signal mitochondrial dysfunction to the integrated stress response.

To express DELE1-mClover at lower levels that would enable its characterization without triggering the ISR, we stably integrated the DELE1-mClover construct at the AAVS-1 safe harbor locus in both wild type and DELE1 knockdown cell lines. This construct was still overexpressed compared to endogenous DELE1, but to a lesser degree than the transiently transfected construct (Extended Data Fig. 5). Expression of DELE1-mClover from the AAVS1 locus was not sufficient to activate the ATF4 reporter in the absence of stress (Fig. 2h), but rescued ISR activation in response to oligomycin in DELE1 knockdown cells (Fig. 2h), demonstrating that the mClover tag does not interfere with this function of DELE1. With this transgene, we confirmed that DELE1 co-localizes with mitochondria (Fig. 2i).

### A cleaved form of DELE1 accumulates upon mitochondrial stress in an OMA1-dependent manner

Characterization of DELE1-mClover by immunoblotting revealed that there are two forms of DELE1, which we named DELE1_L_ and DELE1_S_. The shorter form, DELE1_S_, accumulates after treatment with oligomycin (Fig. 3a) and other mitochondrial toxins (Extended Data Fig. 6). Because the C-terminally mClover-tagged construct is based on DELE1 cDNA, the two observed forms cannot represent alternatively spliced isoforms, but are likely the products of an N-terminal cleavage process. To test this possibility, we subjected purified DELE1_S_ to mass spectrometry. In comparison with total DELE1, three N-terminal peptides were not detected in DELE1s (Fig. 3b). Based on the pattern of detected peptides, we predicted a cleavage site between amino acids 101 and 141. To narrow down the cleavage site, we created truncation constructs of DELE1, and found that DELE1_S_ migrated more closely to a construct lacking the N-terminal 148 amino acids than a construct lacking the N-terminal 100 amino acids (Fig. 3c), suggesting likely cleavage closer to amino acid 141. To identify a potential cleavage motif, we introduced a series of short consecutive amino acid deletions between amino acids 73 and 149. None of these deletions abrogated DELE1 cleavage (Fig. 3d), indicating that there may not be a specific sequence motif dictating the cleavage site, but that cleavage is rather based on the position within the protein. Supporting this hypothesis is the fact that minor bands of different sizes are visible both for full-length DELE1 and for deletion constructs (Fig. 3d), indicating plausible cleavage events at more than one site. However, mitochondrial localization seemed to be required for DELE1 cleavage, since the two N-terminal truncation constructs, ΔN100N and ΔN148, no longer localized to mitochondria (Fig. 3e) and are not cleaved (Fig. 3c).

To identify the potential protease(s) responsible for the cleavage of DELE1, we ranked the knockdown phenotypes of all mitochondrial proteases in our CRISPRi screen. We reasoned that knockdown of the protease would no longer cleave DELE1, and thus reduce ATF4 reporter activation upon mitochondrial stress. Of 33 mitochondrial proteases targeted in our screen, OMA1 knockdown showed the strongest abrogation of ATF4 activation (Fig. 3f). OMA1 is a mitochondrial stress-activated protease localized in the inner mitochondrial membrane^28,29^. We cloned 2 sgRNAs targeting OMA1 and validated their knockdown efficiency via Western blot (Fig. 3g). In OMA1 knockdown cells, DELE1_S_ is reduced to barely detectable levels and ATF4 up-regulation is abolished (Fig. 3g), suggesting that OMA1 cleaves DELE1 and that this cleavage event is necessary to induce ATF4 up-regulation. Conversely, knockdown of the mitochondrial protease IMMP2L, which showed a phenotype in our CRISPRi screen (Fig. 3f), did not interfere with DELE1 cleavage (Extended Data Fig. 7).

Since OMA1 and its known substrate OPA1 localize to the inner mitochondrial membrane^28,29^, we next investigated the submitochondrial localization of DELE1. Two-color, three-dimensional super-resolution microscopy revealed that DELE1 is localized internally with respect to the outer mitochondrial membrane protein Tom20, but externally with respect to the mitochondrial matrix protein Hsp60 (Fig. 3h, Extended Data Fig. 8a), suggesting that DELE1 localizes to the inter-membrane space or the inner mitochondrial membrane. In either location, it would be positioned to physically interact with OMA1. By biochemical fractionation, we found DELE1 to be associated with the mitochondrial membrane fraction (Extended Data Fig. 8b). However, it is challenging to differentiate between peripheral and integral membrane proteins of the inner mitochondrial membrane using classical biochemical approaches^30^.

In summary, we found that a diverse array of mitochondrial stressors stimulates cleavage of DELE1. Given the requirement for both OMA1 and DELE1 for ISR activation upon mitochondrial stress, these findings suggest that DELE1 cleavage products are key mediators in this pathway (Fig. 3i).

### Cytosolic DELE1 physically interacts with and activates HRI

HRI is a cytosolic kinase (based on microscopy, Extended Data Fig. 9b, and biochemical fractionation^31,32^) and we hypothesized that the DELE1_S_ accumulates in the cytosol and interacts with HRI. Indeed, biochemical fractionation indicated that whereas DELE1_L_ is mitochondrially localized, DELE1_S_ accumulates in the cytosol upon mitochondrial stress (Fig. 4a). This could either indicate that mitochondrially localized DELE1 is cleaved and that the resulting DELE1_S_ exits the mitochondria, or that newly synthesized DELE1 is cleaved during mitochondrial import. To distinguish between these possible mechanisms, we treated cells with cycloheximide to block new synthesis of DELE1 during mitochondrial stress. Cycloheximide treatment did not abrogate stress-induced DELE1_S_ accumulation in the cytosol (Fig. 4b), suggesting that cytosolic DELE1_S_ originates from cleavage of pre-existing mitochondrial DELE1_L_. Super-resolution microscopy confirmed an increase of DELE1-mClover signal outside the mitochondria upon oligomycin treatment (Fig. 4c).

To dissect the function of different domains of DELE1 in determining its localization and ability to induce the ISR, we took advantage of the fact that transient overexpression of DELE1 induces ATF4 activation, which enabled systematic structure-function analyses (Extended Data Fig. 9). The C-terminal portion of DELE1 contains 7 predicted TPR repeats, which can mediate protein-protein interactions. Interestingly, a construct containing all TPR repeats but lacking the N-terminal 206 amino acids no longer localized to the mitochondria (Fig. 4d), but still activated the ATF4 reporter when overexpressed (Fig. 4e). This finding indicated that under overexpression conditions, DELE1 does not need to localize to mitochondria to act as an inducer of the ISR. However, removal of N-terminal TPR repeats abrogated DELE1 activity (Fig. 4e, Extended Data Fig. 9). We found that a construct consisting of the first four TPR repeats is the minimal domain required to activate the ATF4 reporter (Extended Data Fig. 9), suggesting that these repeats mediate a protein-protein interaction necessary for the activation of the ISR.

Next, we asked whether HRI and DELE1 interact with each other in the cytosol. Indeed, HRI interacted with DELE1 based on co-immunoprecipitation (Fig. 4f). This interaction was dependent on the N-terminal TPR repeats (Fig. 4f), mirroring the requirement for ATF4 activation.

To test if the interaction of DELE1 with HRI is sufficient to activate the eIF2α kinase activity of HRI, we assayed HRI kinase activity *in vitro* using purified HRI, eIF2α and DELE1 (Fig. 4g). Indeed, stoichiometric amounts of DELE1 stimulated HRI-mediated eIF2α phosphorylation, both in the presence and absence of hemin (Fig. 4h). Thus, we identified a novel mechanism of HRI activation that operates even in the presence of high heme concentrations. Taken together, our results suggest that the stress-induced accumulation of DELE1 in the cytosol leads to the activation of HRI through physical interaction (Fig. 4i).

### The DELE1-HRI pathway can be maladaptive, and its blockade induces an alternative response

Depending on its context and duration, the ISR can have either homeostatic or pro-apoptotic outcomes, and it can be deficient or maladaptive in disease states^17,33^. We therefore asked whether the DELE1-HRI-mediated response to mitochondrial stress is beneficial to cells. Treatment of HEK293T cells with 2.5 ng/ml oligomycin for 16 hours reduced the number of live cells; this reduction was abolished by HRI knockdown (Fig. 5a), indicating a maladaptive function of the ISR in this context. Using different genetic perturbations of mitochondrial functions, we found that HRI knockdown was similarly beneficial to cells with a depleted mitochondrial ribosomal protein (Fig. 5b). Conversely, HRI knockdown sensitized cells to knockdown of the mitochondrial protease LonP1 (Fig. 5b), indicating that the ISR plays a protective role in LonP1 knockdown cells.

**Fig. 5.**
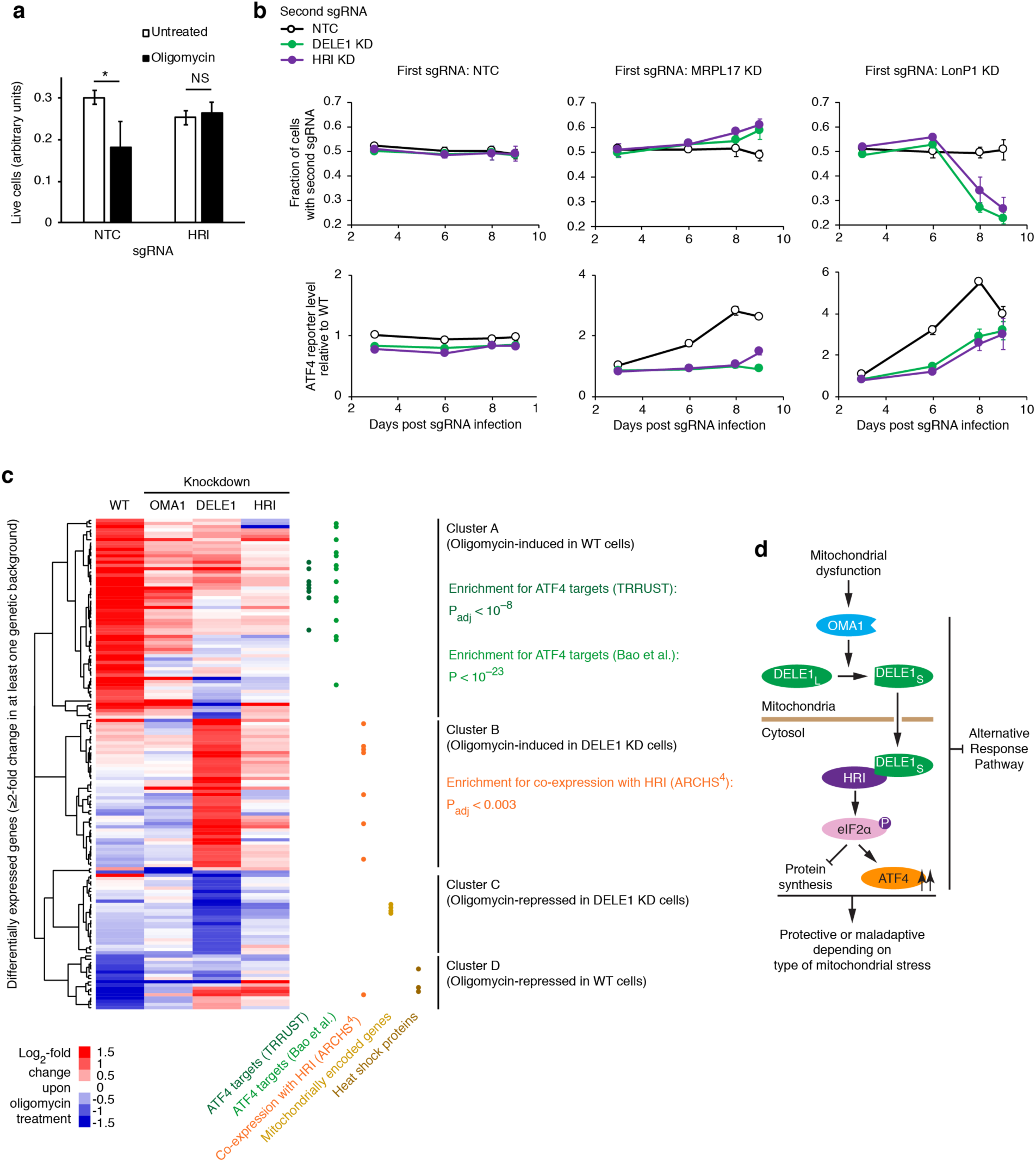
The DELE1-HRI pathway can be maladaptive and its blockade induces an alternative program. **(a)** HRI knockdown is protective during oligomycin treatment. HEK293T cells expressing non-targeting control sgRNA (NTC) or an sgRNA knocking down HRI were untreated or treated with 2.5 ng/mL oligomycin for 16 h, and cell numbers were determined by counting (mean ± s.d., *n* = 3 culture wells, \*\**P* < 0.05, two-tailed unpaired *t* test). **(b)** HRI knockdown is protective for cells with depleted mitochondrial ribosomal protein MRPL17, but sensitizes cells with depleted mitochondrial protease LonP1. HEK293T cells were co-infected with lentiviral construct expressing green fluorescent protein and an sgRNA knocking down HRI, and with a lentiviral construct expressing blue fluorescent protein and a non-targeting control sgRNA (NTC) or an sgRNA knocking down MRPL17, MRPL22, or LONP1. Cells were cultured for 9 days and proportions of cells expressing green and blue fluorescent proteins were quantified on days 3, 6, 8 and 9 post infection by flow cytometry (*top*). Thus, the effect of HRI knockdown on proliferation in different genetic backgrounds could be evaluated in an internally controlled experiment. In parallel, the ATF4 reporter was quantified (*bottom*). Mean ± s.d., *n* = 3 culture wells. **(c)** HEK293T cells that were either infected with a non-targeting control sgRNA or in which OMA1, DELE1 or HRI was knocked down were untreated or treated with 1.25 ng/mL oligomycin for 16 h, and differentially expressed genes were detected by RNA sequencing. The heatmap only includes genes expression of which changed significantly upon oligomycin treatment (p_adj_ < 0.05) by at least two-fold in at least one genetic background. (Full datasets are provided as Supplemental Tables 2, 4-6). Hierarchical clustering reveals four major gene clusters. Gene groups are indicated by dots in different colors: ATF4 targets annotated by the TRRUST database (dark green) or Bao *et al*.^11^ (light green), genes co-expressed with HRI in the ARCHS^4^ database (orange dots) mitochondrially encoded genes (golden dots), heat-shock proteins (brown dots). **(d)** Final model.

To characterize the response to mitochondrial stress in cells lacking the OMA1-DELE1-HRI pathway, we conducted RNA-Seq in different genetic backgrounds and compared genes that were differentially expressed upon oligomycin treatment (Fig. 5c, Extended Data Fig. 10). Hierarchical clustering revealed four major gene clusters. Genes that are induced by oligomycin in WT cells (Cluster A) are induced to a substantially lesser degree in OMA1, DELE1 and HRI knockdown cells. These genes are enriched for ATF4 targets annotated by the TRRUST database^34^, in which ATF4 was the only significantly enriched transcription factor (p_adj_ < 0.05) for Cluster A genes, and Bao *et al*.^11^. This finding confirmed that knockdown of the newly identified pathway not only reduced the translational induction of ATF4, but also reduced the transcriptional induction of ATF4 target genes.

Clusters B-D were not significantly enriched (p_adj_ < 0.05) for targets of any transcription factor in TRRUST. However, genes induced by oligomycin in DELE1 knockdown cells (ClusterB) were enriched for co-expression with HRI in the ARCHS^4^ database^35^ – and HRI was the only significantly enriched kinase for co-expression with Cluster B genes. Cluster C (genes repressed by oligomycin in DELE1 knockdown cells) contained four mitochondrially encoded genes, possibly reflecting transcriptional regulation of the mitochondrial genome, or loss of mitochondrial RNA through mitophagy. Cluster D (genes repressed by oligomycin in WT cells) contained two cytosolic Hsp70 heat-shock proteins. Upon DELE1 or HRI knockdown, the inducible Hsp70 (HSPA1B) is induced, rather than repressed, in response to oligomycin. This last finding is particularly intriguing, since it indicates that a blockade of the ISR in the context of mitochondrial stress can lead to upregulation, instead of downregulation, of Hsp70, which may mediate in part the protective effect of HRI knockdown we observed in some stress contexts (Fig. 5a,b).

In summary, the OMA1-DELE1-HRI pathway can be protective or maladaptive, depending on the specific type of mitochondrial stress cells experience, and its blockade leads to the induction of an alternative stress response pathway (Fig. 5d).

### Concluding remarks

Using an unbiased genetic screen, we identified the molecular players (OMA1, DELE1, HRI) in a pathway that signals mitochondrial dysfunction to the ISR. We thereby provide a novel cellular role and mechanism of activation for HRI. In conjunction with the recently reported role of HRI in innate immune signaling and its regulation by the heat shock protein HSPB8^36^, we thus further expand our understanding of HRI as a regulator beyond its canonical role as a heme sensor.

Our findings also raise new questions that will be the subject of future studies.

How does OMA1 sense mitochondrial stress to trigger DELE1/HRI mediated activation of the ISR? Is the underlying regulatory mechanism the same as for stress-induced cleavage of OPA1 by OMA1, or do distinct mechanisms control DELE1 cleavage?

How does DELE1_S_ get transported to the cytosol? Does this mechanism resemble that of PGAM5, which has similarly been reported to be cleaved at the inner mitochondrial membrane upon stress, followed by the release of its C-terminal fragment into the cytosol^37^?

Given that some DELE1_S_ is present in the cytosol in the absence of mitochondrial stress, are there additional layers of regulation? These may involve post-translational modifications of DELE1 or HRI, and additional protein interaction partners. It is tempting to speculate that Hsp70 and/or Hsp90 chaperones may play a role, given their reported modulation of HRI activity^20,38,39^ and the fact that DELE1 contains a TPR domain, which in some proteins mediates physical interactions with Hsp70 and Hsp90.

Is there cross-talk with other regulatory pathways? While our work provides strong evidence for HRI as the major eIF2α kinase mediating ISR activation in response to mitochondrial stress, other eIF2α kinases may contribute once the cellular stress becomes more generalized. Previous studies have implicated GCN2^40^ and PERK^41^, and our own results suggest that even DELE1 and HRI knockdown cells can activate the ISR in a delayed manner after several days of mitochondrial stress (Fig. 5b, LonP1 knockdown cells). Translation of ATF4 also requires basal mTOR activity^42,43^, but there is currently no direct evidence supporting that mTOR activation is the signal driving ISR activation in response to mitochondrial stress.

Blockade of the OMA1-DELE1-HRI pathway in the presence of mitochondrial stress surprisingly led to the activation of an alternative response pathway, which may account for the beneficial effects of DELE1 and HRI knockdown in the context of some, but not all mitochondrial stresses (Fig. 5ab). The molecular mechanism controlling this alternative pathway remains to be elucidated.

The central role of the ISR has generated a lot of interest in its potential role as a therapeutic target. Both pharmacological inhibition^44^ and prolongation^45^ of the ISR have shown promise in animal models of disease. However, the context-dependent function of the ISR, which can be protective or maladaptive, presents a challenge. Indeed, our finding that ISR blockade during mitochondrial dysfunction leads to activation of an alternative response highlights the complexity of targeting the ISR.

The OMA1-DELE1-HRI pathway we describe here is a potential therapeutic target for blocking ISR activation in cells experiencing mitochondrial dysfunction, without globally blocking the ISR in all cells. OMA1 ablation has previously been reported to protect against heart failure in multiple mouse models that involve mitochondrial dysfunction^46^. Our results suggest that this effect may be mediated by attenuation of ISR activation. DELE1 could be an even more attractive therapeutic target, since OMA1 performs independent homeostatic functions, and since DELE1 has an additional pro-apoptotic activity^27^. Furthermore, DELE1 knockdown resulted in the most stringent inhibition of ISR activation and induction of the alternative stress response pathway (Fig. 5c).

By dissecting the molecular pathways controlling the ISR under different stress conditions, additional therapeutic targets may emerge, which could enable fine-tuned manipulation of the cellular response to different stressors to achieve beneficial outcomes tailored to specific disease states.

## Acknowledgments

We thank Greg Mohl and Ben Herken for contributions to preliminary experiments, Jaime Leong for FACS support, Christopher Richards for contributions to data visualization, Kevan Shokat and members of the Kampmann lab for discussions, Adam Frost, Isha Jain, Avi Samelson and Emmy Li for comments on the manuscript, Eric Chow (UCSF Center for Advanced Technology) for support with next-generation sequencing, Sarah Elmes (UCSF Laboratory for Cell Analysis) for support with FACS, Delaine Larson (UCSF Nikon Imaging Center) for support with fluorescence microscopy. This work was supported by the National Institutes of Health grants GM119139 (M.K.), DK26506 (M.A.C.), GM44037 (M.A.C.), OD022552 (A.P.W.), the Beckman Young Investigator Program (K.X.) and a Larry L. Hillblom Foundation Postdoctoral Fellowship (X.G.). M.K. and K.X. are Chan Zuckerberg Biohub Investigators.

## Author contributions

X.G. and M.K. conceptualized and led the overall project, analyzed results and wrote the manuscript, with input from all co-authors. X.G. and G.A. conducted and analyzed the CRISPRi screen and follow-up experiments. Y.L. and M.A.C. conducted and analyzed experiments with purified proteins. R.T. and M.K. analyzed and visualized RNA-Seq results. B.U. and K.X. conducted and analyzed super-resolution experiments. Y.T.L and A.P.W. conducted and analyzed mass spectrometry experiments.

## Author information

The authors declare no competing interests. Correspondence and requests for materials should be addressed to.

## METHODS

### Plasmids

Sequences of oligonucleotides used for cloning are provided in Extended Data Table 1. The ATF4 translational reporter (pXG237) was generated by replacing the venus-IRES-BFP of pMK1163, our previously published integrated stress response vector^47^, with mApple through Gibson assembly (NEB, E2611). The secondary reporter (pXG260) to control the transcriptional regulation of CMV was generated by replacing ATF4-uORFs1/2-mApple with EGFP through Gibson assembly.

The DELE1 coding sequence was cloned from human cDNA and inserted into pMK1253^48^ through Gibson assembly to generate a DELE1-mClover tagged protein under the EIF1A promoter (pXG286). To obtain a construct that enabled integrate DELE1-mClover at the safe harbor locus (pXG289), the EIF1A promoter-DELE1-mClover cassette was PCR-amplified from pXG286, and inserted between the SpeI and MluI sites of the AAVS-1-targeting vector pMTL3^26^. All truncation constructs were generated through ligation of the corresponding truncated DELE1 into EcoRI- and NotI-digested pXG286. Internal short consecutive deletions covering amino acids 73 to 149 were made by inverse PCR on pXG286.

HRI cDNA was synthesized as a gene block by IDT and cloned into pMK1253 as for pXG286 to obtain pXG272.

To clone individual sgRNAs, top and bottom oligonucleotides (IDT) were annealed and ligated to our optimized lentiviral sgRNA expression vector^19^. Triple sgRNAs expression constructs were generated as previously published^49^. Protospacer sequences for sgRNAs are listed in Extended Data Table 2.

### Cell lines

HEK293T cells were cultured in DMEM (Gibco, 11965-092) with 10% fetal bovine serum (Seradigm # 97068-085, Lot# 076B16), Pen/Strep (Life Technologies # 15140122), and L-glutamine (Life Technologies # 25030081).

The CRISPRi HEK293T cell line (cXG284) was generated by transfecting HEK293T cells with pC13N-dCas9-BFP-KRAB^26^ and TALENS targeting the human CLYBL intragenic safe harbor locus (between exons 2 and 3) (pZT-C13-R1 and pZT-C13-L1, Addgene #62196, #62197) using DNA-In Stem (VitaScientific). BFP-positive cells were isolated via FACS sorting.

The ATF4 translational reporter cell line (cXG289) was generated through lentiviral infection of cXG284 with pXG237 and FACS-based monoclonal selection based on response to mitochondrial stress. The dual reporter cell line (cXG330) was generated via lentiviral infection of the second reporter (pXG260) into cXG289.

CRISPRi knockdown cell lines were generated by lentiviral transduction with plasmids containing individual sgRNAs or triple sgRNAs.

The cell line expressing a DELE1-mClover transgene line from the AAVS1 locus was generated by transfecting a cXG289 population in which ∼ 50% of cells expressed a DELE1-sgRNA and a BFP marker with pXG289 and TALENS targeting the human AAVS1 locus (AAVS1-TALEN_L/R, Addgene #59025 and #59026). Through FACS sorting, cells expressing the DELE1-mClover transgene either in the wild type background (BFP-, GFP+) or in the DELE1 knockdown background (BFP+, GFP+) were isolated.

### Drug treatments

HEK293T cells and derived cell lines were seeded at 25% confluency 24 h before drug treatment. An equal volume of DMEM with twice the final drug concentration was added. To trigger ER stress, cells were treated with 75 nM of thapsigargin (Sigma-Aldrich # T9033) for 8 h. For mitochondrial stress, cells were incubated with the following mitochondrial toxins for 16 h: 1.25 ng/mL oligomycin (Sigma-Aldrich # 75351), 50 µg/mL doxycycline (Clontech # 631311), 40 nM Antimycin (Sigma-Aldrich # A8674), 40 nM Rotenone (Sigma-Aldrich # R8875) and 5 µM CCCP (Sigma-Aldrich # C2759). For hemin supplementation experiments, 10 μM or 20 μM hemin (Sigma-Aldrich # H9039) was added to the medium when cells were seeded before oligomycin treatment, or hemin was only added during oligomycin treatment. HEK293 cells were treated with 10 μM or 20 μM for 24 h and harvested to quantify mRNA levels of HO-1. For cycloheximide (CHX) (Sigma-Aldrich # A4859) experiments, cells were treated with 20 μg/mL of CHX for 4 h.

### Western blot

Cells were lysed using RIPA buffer (Thermo Fisher Scientific #89900). Total protein was quantified via Pierce BCA Protein Assay Kit (Thermo Fisher Scientific #23225). Samples were subjected to SDS-PAGE on NuPage 4-12% Bis-tris gels Bis-tris gels (Thermo Fisher Scientific # NP0336BOX) and transferred to nitrocellulose membrane (Bio-Rad #1704271). The primary antibodies used in this study were: rabbit anti-ATF4 (Cell Signaling Technologies #11815, 1:500) mouse anti-β-actin (Cell Signaling Technologies #3700, 1:5000), mouse anti-GFP (Roche #11814460001, 1:1000), rabbit anti-HRI (Mybiosource #MBS2538144, 1/1000), rabbit anti-OMA1 (Cell Signaling Technologies #95473, 1:1000), rabbit anti-LonP1 (Cell Signaling Technologies #56266, 1:1000), rabbit anti-Hsp60 (Cell Signaling Technologies #12165, 1:1000), mouse OXPHOS cocktail (anti-ATP5A, anti-UQCRC2, anti-SDHB, anti-COX II and anti-NDUFB8) (Abcam #ab110411, 1:1000), rabbit anti-AIF (Cell Signaling Technologies #5318, 1:1000). Antibodies failed to detect DELE1 include: Abcam #ab189958, 1:500; Santa Cruz Biotech # sc-515080, 1:100; Proteintech # 21904-1-AP, 1:500, Biorbyt # ABIN1031350, and 1:500. Blots were incubated with Li-Cor secondary antibodies and imaged via Odyssey Fc Imaging system (Li-Cor #2800). Digital images were processed and analyzed using Licor ImageStudio™software.

### Quantitative RT-PCR

Total RNA was extracted using the Quick-RNA Miniprep Kit (Zymo Research #R1054), and first strand cDNA was synthesized with the SuperScript III First-Strand Synthesis System (Invitrogen #18080-051). qPCR was performed with SensiFAST SYBR Lo-ROX reagents (Bioline #BIO-94005). Expression fold changes were calculated using the ΔΔCt method. qPCR primers used are listed in Extended Data Table 3.

### CRISPRi screen

To obtain pooled sgRNA virus, next-generation sgRNA libraries H1, H3 and H4^24^ were transfected into HEK293T cells together with lentiviral plasmid packaging mix using TransIT^®^-Lenti Transfection Reagent (Mirus #MIR 6600). The dual reporter cell line (cXG330) was then transduced with the pooled sgRNA virus and selected with puromycin (2.5 μg/mL) for 2-3 days until greater than 90% of the cells were BFP-positive. Cells were then cultured for 3 days in the absence of puromycin. Cell populations were cultured such that a representation of at least 1000 cells per sgRNA element was maintained throughout the screen. Cells were seeded at 2.5 million per 10 cm dish (12 dishes total) at a volume of 7.5 mL DMEM on day 0. On day 1, 7.5 mL of additional DMEM with or without 2.5 ng/mL oligomycin was added into each dish. On day 2, after 16 h of oligomycin treatment, both untreated and oligomycin-treated cells were FACS-sorted based on the ratio of mApple to GFP fluorescence intensity, and population corresponding to the top 30% and bottom 30% of cells were collected. This experimental design is optimal for FACS-based screen based on our previous simulations^50^. The representation in each sorted population was ∼500 cells per sgRNA element. Genomic DNA was isolated using a Macherey-Nagel Blood L kit (Machery-Nagel #740954.20). sgRNA-encoding regions were amplified and sequenced as previously published^19^. Phenotype and P value for each gene were calculated using our recently described bioinformatics pipeline^26^. Gene scores were defined as the product between the phenotype and -log_10_(P value). Full screen results are provided as Supplemental Table 3.

### Cellular fractionation

Cytosolic and mitochondrial fractions were separated using the Mitochondrial Isolation Kit for Mammalian Cells (Thermo Fisher Scientific, #89874). To separate submitochondrial fractions, isolated mitochondria were processed with the Mem-PER™ plus kit (Thermo Fisher Scientific, #89842). To extract peripheral membrane proteins, 0.1 M Na_2_CO_3_ (pH = 11) were added for a 30 min incubation at 4 °C between the permeabilization and solubilization steps.

### Immunoprecipitation

HEK293T cells were transiently transfected with DELE1-mClover (pXG286) or DELE1(Δ275)-mClover constructs. Cells were collected 24 h after transfection and lysed using a mild lysis buffer (10 mM Tris/Cl pH 7.5; 150 mM NaCl; 0.5 mM EDTA; 0.5% NP-40, 0.09% Na-Azide). The lysates were then incubated with GFP-Trap®_MA beads (Chromtek, #gtma20) for one hour at 4°C. Proteins captured on the magnetic beads were boiled in 2X SDS loading dye for 10 min before subjecting to SDS PAGE and Western blotting.

### Confocal microscopy

HEK293T cells and derived cell lines were seeded in the 8-well chamber slide (Ibidi #80824) for 24 h and transfected with DELE1 truncated constructs and/or plasmid mito7-mRuby (a gift from Michael Davidson, Addgene plasmid # 55874) as indicated, using TransIT-Lenti Transfection Reagent (Mirus #MIR 6600). 24 h after transfection, cells were incubated with 100 nM MitoTrackerTM red CMXRos (ThermoFisher Scientific #M7512) for 20 min at 37°C where indicated. Live cell images were then taken using the Yokagawa CSU22 spinning disk confocal and processed using Fiji^51^.

### Super-resolution microscopy

#### Cell fixation and immunofluorescence

DELE1-mClover cells were seeded on #1.5 glass coverslips and cultured for 24 h, followed by treatment with 1.25 ng/ml oligomycin for 16 h where indicated. Samples were fixed with 3% (w/v) paraformaldehyde and 0.1% (w/v) glutaraldehyde in phosphate-buffered saline (PBS) for 20 min. After reduction with a freshly prepared 0.1% sodium borohydride solution in PBS for 5 min, the samples were permeabilized and blocked in a blocking buffer (3% w/v BSA,%0.1 % v/v Triton X-100 in PBS) for 1 h. Afterward, the cells were incubated with primary antibodies (below) in the blocking buffer for 12 h at 4 °C. After washing in a washing buffer (0.3% w/v BSA and 0.01% v/v Triton X-100 in PBS) for three times, the cells were incubated with dye-labeled secondary antibodies (below) for 1 h at room temperature. Then, the samples were washed 3 times with the washing buffer and 3 times with PBS. Primary antibodies used: mouse anti-mClover/GFP (Invitrogen #A-11120, 1:350); rabbit anti-Tom20 (Santa Cruz Biotechnology #sc-11415, 1:100); rabbit anti-Hsp60 (Cell Signaling Technology #12165, 1:800). Secondary antibodies used: Alexa Fluor 647-labeled goat anti-mouse (Invitrogen #A21236, 1:400); donkey anti-rabbit (Jackson ImmunoResearch #711-005-152, 1:70) conjugated with CF568 succinimidyl ester (Biotium #92131).

#### Super-resolution microscopy

3D-STORM super-resolution microscopy^52,53^ was carried out on a homebuilt setup using a Nikon CFI Plan Apo λ 100x oil immersion objective (NA 1.45), as described previously^54^. Briefly, the sample was mounted with an imaging buffer consisting of 5% (w/v) glucose, 100 mM cysteamine, 0.8 mg/mL glucose oxidase, and 40 µg/mL catalase in a Tris-HCl buffer (pH 7.5).The labeled Alexa Fluor 647 and CF568 molecules were sequentially photoswitched to the dark state and imaged as single molecules using 647- and 560-nm lasers; these lasers were passed through an acousto-optic tunable filter and illuminated a few micrometers into the sample at ∼2 kW cm^-2^. Single-molecule emission was passed through a cylindrical lens of focal length 1 m to introduce astigmatism^53^, and recorded with an Andor iXon Ultra 897 EM-CCD camera at a framerate of 110 Hz, for a total of ∼50,000 frames per image. Data acquisition used publicly available software (https://github.com/ZhuangLab/storm-control). The raw STORM data were analyzed using Insight3 software^53^ according to previously described methods^52,53^.

### Mass spectrometry

#### Sample Preparation for Mass Spectrometry

DELE1-mClover was affinity purified from mitochondrial and cytosolic fractions of cells with DELE1-mClover in the AAVS1 locus using GFP-Trap®_MA beads (Chromtek #gtma20). Beads and affinity-purified proteins were resuspended in a buffer containing 4M guanidine hydrochloride, 0.1 M Tris pH 8.0, 5 mM tris(2-carboxyethyl)phosphine (Sigma #C4706), and 10 mM 2-chloroacetamide (Sigma #22790). The mixture was incubated at room temperature for 1 hour for protein reduction and alkylation. Subsequently, 1 M Tris pH 8.0 was used to dilute the concentration of guanidine hydrochloride to 1M followed by the addition of 5 μg of mass spectrometry-grade trypsin (Thermo Fisher Scientific #90057). The digestion reaction was incubated at 37°C for 24 h on a shaker. Soluble tryptic peptides were collected by bead separation on a magnetic stand. Peptides were acidified to a final concentration of 1% trifluoroacetic acid (pH < 3), desalted by SOLA C18 solid phase extraction (SPE) cartridge (Thermo Fisher Scientific #60109-001), and then dried down in a speed-vac. Dried peptides were stored at −80°C until mass spectrometry analysis.

#### Liquid Chromatography-Tandem Mass Spectrometry (LC-MS/MS)

Desalted peptides were re-constituted in 2% acetonitrile, 0.1% formic acid and diluted to 0.2μg/μL before mass spectrometry analysis. For each sample, a total of 1 ug of peptides was injected into a Dionex Ultimate 3000 NanoRSLC instrument with a 15-cm Acclaim PEPMAPC18 (Thermo Fisher Scientific #164534) reverse phase column. The samples were separated on a 2-hour non-linear gradient using a mixture of Buffer A (0.1% FA) and B (80% ACN/0.1% FA). The initial flow rate was 0.5 uL/min at 3% B for 13 minutes followed by a drop in flow rate to 0.2 uL/min and a non-linear increase (curve 7) to 40% B for the next 83 minutes. The flow rate was then increased to 0.5 uL/min while Buffer B was linearly ramped up to 99% for the next two minutes. Finally, we maintained the peak flow rate and Buffer B concentration for another five minutes before lowering Buffer B to 3%. Eluted peptides were analyzed with a Thermo Q-Exactive Plus mass spectrometer. The MS survey scan was performed over a mass range of 350-1500 m/z with a resolution of 70,000. The automatic gain control (AGC) was set to 3e6, and the maximum injection time (MIT) was 100 ms. We performed a data-dependent MS2 acquisition at a resolution of 17,500, AGC of 5e4, and MIT of 150 ms. The 15 most intense precursor ions were fragmented in the HCD at a normalized collision energy of 27. Dynamic exclusion was set to 20 seconds to avoid over-sampling of highly abundant species.

#### MS Data Processing

We analyzed the raw spectral data using MaxQuant version 1.5.1.2^55^ for protein identification by searching against the DELE1 sequence (Q14154 UniProtID). We used the “peptides” output file for downstream analyses.

### Recombinant proteins

Full length N-terminal (His)_6_-tagged rat HRI in construct pET28a-(His)_6_-HRI^56^ was co-expressed in *Escherichia coli* BL21(DE3) with chaperone plasmid pG-KJE8 (Takarabio) at 15°C for 72 h. Cleared cell lysates (lysed in Ni column buffer: 50 mM KPB pH 8.0, 300 mM NaCl, 10% glycerol, 5 mM imidazole, 0.5% CHAPS supplemented with protease inhibitor cocktail) were first applied to a His60 Ni Superflow column (Clontech). Eluates from the His60 Ni column were pooled and dialyzed overnight against 50 mM sodium phosphate pH 7.4 and then loaded onto a DEAE column. (His)_6_HRI was then step eluted with increasing concentrations of NaCl. Elution fractions with 200 mM, 250 mM and 300 mM NaCl were pooled and concentrated and then further purified through gel filtration on a HiPrep S-300 column and then concentrated and buffer exchanged into PBS with 20% glycerol using an Amicon centrifugal filter column.Purified (His)_6_-HRI proteins were more than 95% homogeneous as evidenced by SDS-PAGE.

Yeast (His)_6_-eIF2α (1-200) in the pET-15b vector^57^ was expressed in *Escherichia coli* BL21(DE3) at 28°C for 16 h. Protein was purified on a His60 Ni Superflow column (Clontech) according to the manufacturer’s instructions. Briefly, cell pellets were lysed in Ni column buffer and then loaded onto the His60 Ni Superflow column. The column was first washed using lysis buffer supplemented with 20 mM imidazole, and then washed using lysis buffer supplemented with 50 mM imidazole. (His)_6_-eIF2α was eluted using lysis buffer supplemented with 300 mM imidazole. Different elution fractions were collected and aliquots were checked by SDS-PAGE. Highly pure fractions (greater than 95% homogeneous) were combined and concentrated using an Amicon centrifugal filter column and finally stored in PBS with 20% glycerol.

Full-length recombinant human DELE1 protein fused to GST was purchased from Abcam (#ab160664).

### HRI kinase assay

25 nM HRI with or without 5 μM hemin were first incubated with 25 nM (1:1) or 50 nM (1:2) DELE1 in kinase buffer (20 mM Tris buffer pH 7.4, 40 mM KCl, 3 mM Mg acetate, 1 mM DTT) at room temperature for 20 min. The reactions were initiated by adding 0.5 μM eIF2α and 50 μM ATP and then incubated at room temperature. The reactions were terminated by addition of SDS sample buffer. Aliquots were subjected to SDS-PAGE, transferred to nitrocellulose membrane, and then immunoblotting with phospho-eIF2α (Ser52) antibody (Invitrogen, **#**44-728G). Blots were scanned using Licor for quantification.

### RNA sequencing and analyses

RNA was extracted from cells using the Quick-RNA Miniprep Kit (Zymo; Cat. No. R1054) and sent to the DNA Technologies and Expression Analysis Core at the UC Davis Genome Center where they performed 3’ Tag-seq, supported by NIH Shared Instrumentation Grant 1S10OD010786-01.

To obtain transcript abundance counts, sequencing reads from 3’-Tag RNA-seq were mapped to human reference transcriptome (GRCh38, Ensembl Release 97) using Salmon v0.14.1^58^ with the “--noLengthCorrection” option. Gene-level count estimates were obtained using tximport v1.8.0^59^ with default settings. Subsequently, genes with more than 10 counts were retained for differential gene expression analysis using DESeq2 v1.20.0^60^. Result files from DESeq2 analysis are provided as Supplemental Tables 1,2,4-6. Significantly enriched gene sets were determined using Enrichr^61,62^. P values for overlap with ATF4 targets from Bao *et al*. ^11^ were calculated using Fisher’s exact test. Hierarchical clustering of differentially expressed genes was carried out in Cluster 3.0^63^ and results were visualized using Java Tree View^64^.

## Data availability

RNA sequencing data described in this manuscript (associated with Fig. 5c and Extended Data Fig. 1) will be deposited in the NCBI Gene Expression Omnibus. There are no restrictions on data availability.

## Code availability

Analysis of CRISPRi screen results was carried out using custom code, MAGeCK-iNC, developed in the Kampmann lab, which was previously described^26^ and is freely available at https://kampmannlab.ucsf.edu/mageck-inc.

## EXTENDED DATA

**Extended Data Figure 1.**
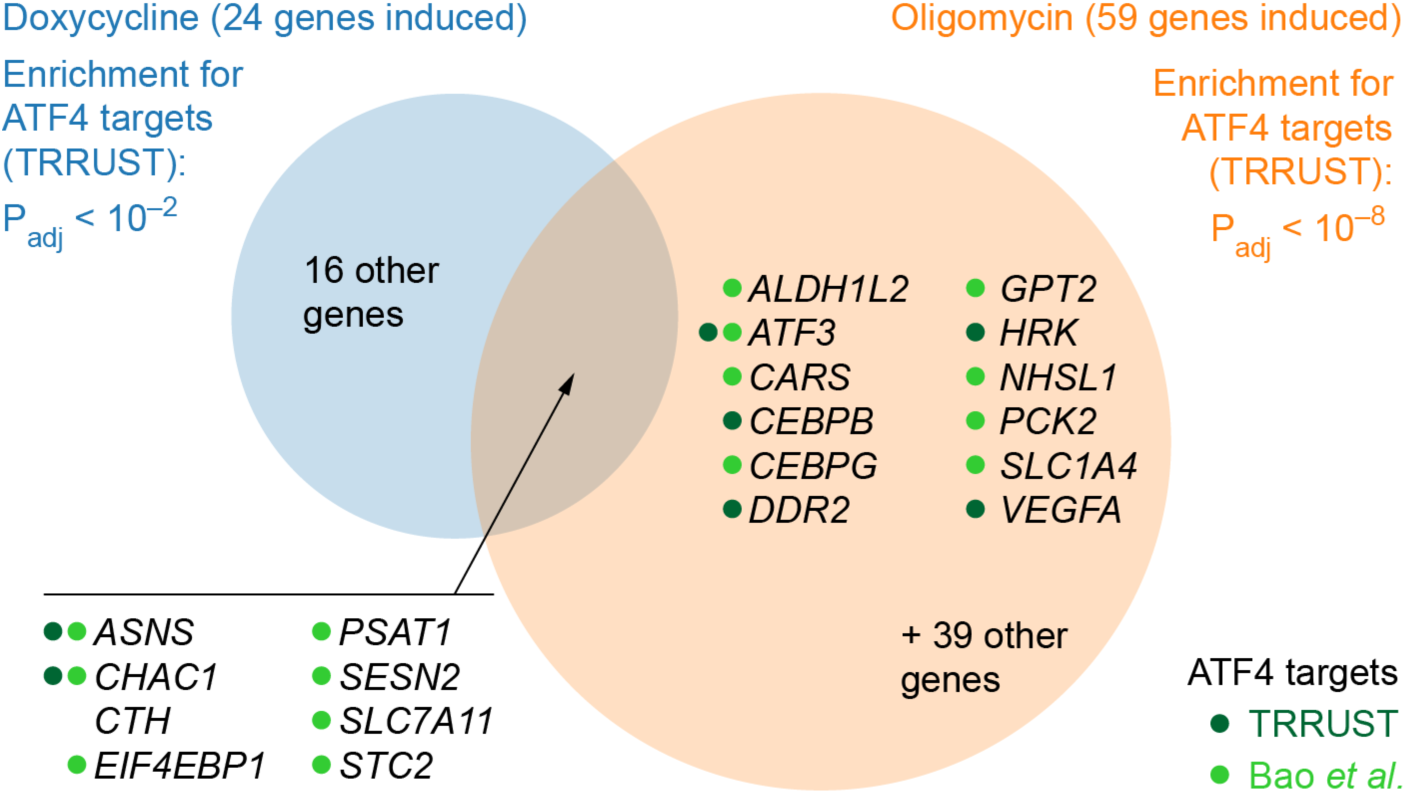
Mitochondrial stressors doxycycline and oligomycin induce ATF4 target genes. HEK293T cells were treated with 50 µg/mL doxycycline or with 1.25 ng/mL of oligomycin for 16 h, and transcript levels were compared to untreated cells using RNA-Seq. Differentially expressed genes were determined (full datasets in Supplemental Tables 1,2). This Figure analyzes significantly induced genes (p_adj_ < 0.05) with at least a 2-fold increase in treated over untreated conditions. Enrichment analysis for targets of transcription factors annotated in TRRUST detected ATF4 as the only significant transcription factor (p_adj_ < 0.05) for both treatments. Genes induced by both treatments are listed, as well as genes annotated as ATF4 targets in TRRUST (dark green dots) or Bao et al. (light green dots).

**Extended Data Figure 2.**
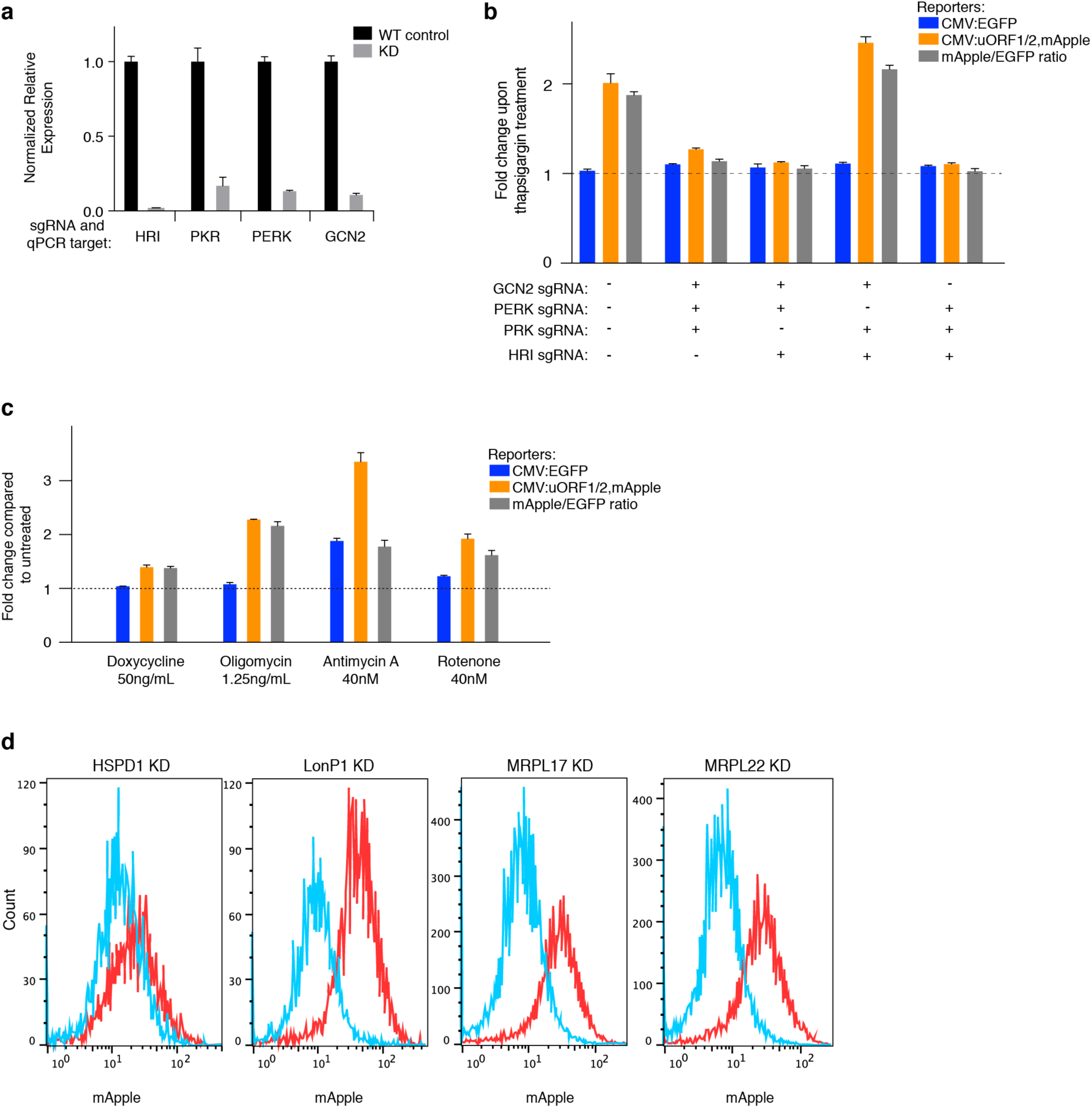
Characterization of the ATF4 translational reporter. **(a)** Quantitative RT-PCR quantification of the knockdown efficiency of four sgRNAs targeting the eIF2α kinases. **(b**) Tharpsigargin induces the ATF4 reporter in a PERK-dependent manner. Reporter cells expressing triple sgRNAs targeting the indicated eIF2α kinases were exposed to 75 nM thapsigargin for 8 h before measuring reporter levels by flow cytometry. The reporter fold change (mean ± s.d., *n* = 3 culture wells) is the ratio of median fluorescence values for thapsigargin over untreated samples. **(c)** Pharmacological inhibition of mitochondrial function induces the ATF4 reporter. Reporter cells were exposed to the indicated treatments for 16 h before measuring reporter levels by flow cytometry. The reporter fold change (mean ± s.d., *n* = 3 culture wells) is the ratio of median fluorescence values for treated over untreated samples. **(d)** CRISPRi knockdown (KD) of genes essential for mitochondrial functions (red) induce the ATF4 reporter compared to WT cells (blue).

**Extended Data Figure 3.**
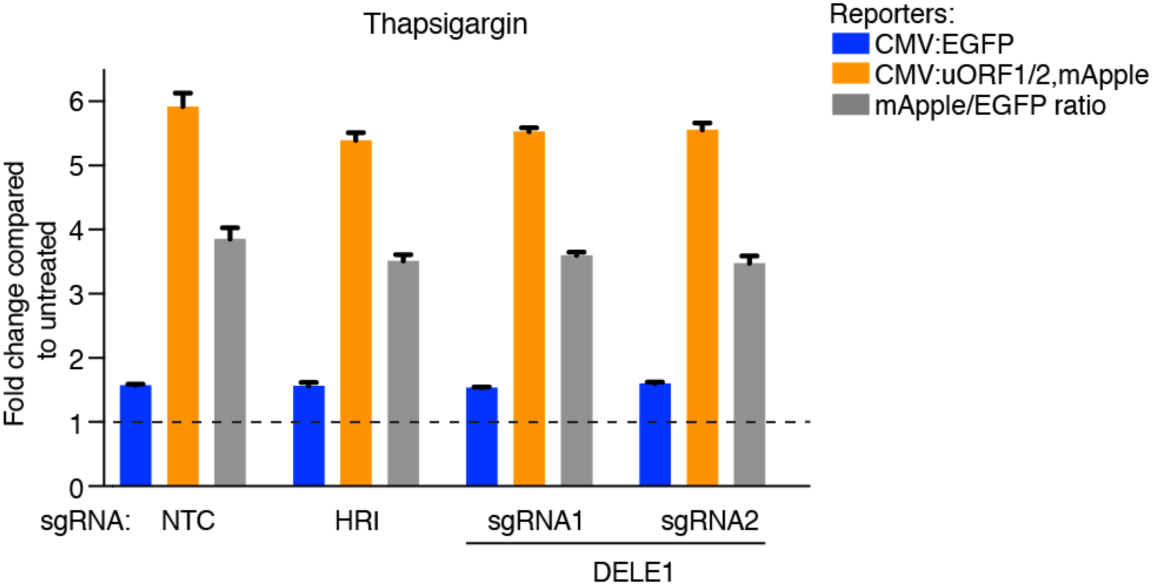
DELE1 and HRI are not required to trigger the integrated stress response in response to ER stress. Reporter cells expressing non-targeting control sgRNAs (NTC) or sgRNAs targeting HRI or DELE1 were exposed to 75 nM thapsigargin for 8 h before measuring reporter levels by flow cytometry. The reporter fold change (mean ± s.d., *n* = 3 culture wells) is the ratio of median fluorescence values for thapsigargin over untreated samples.

**Extended Data Figure 4.**
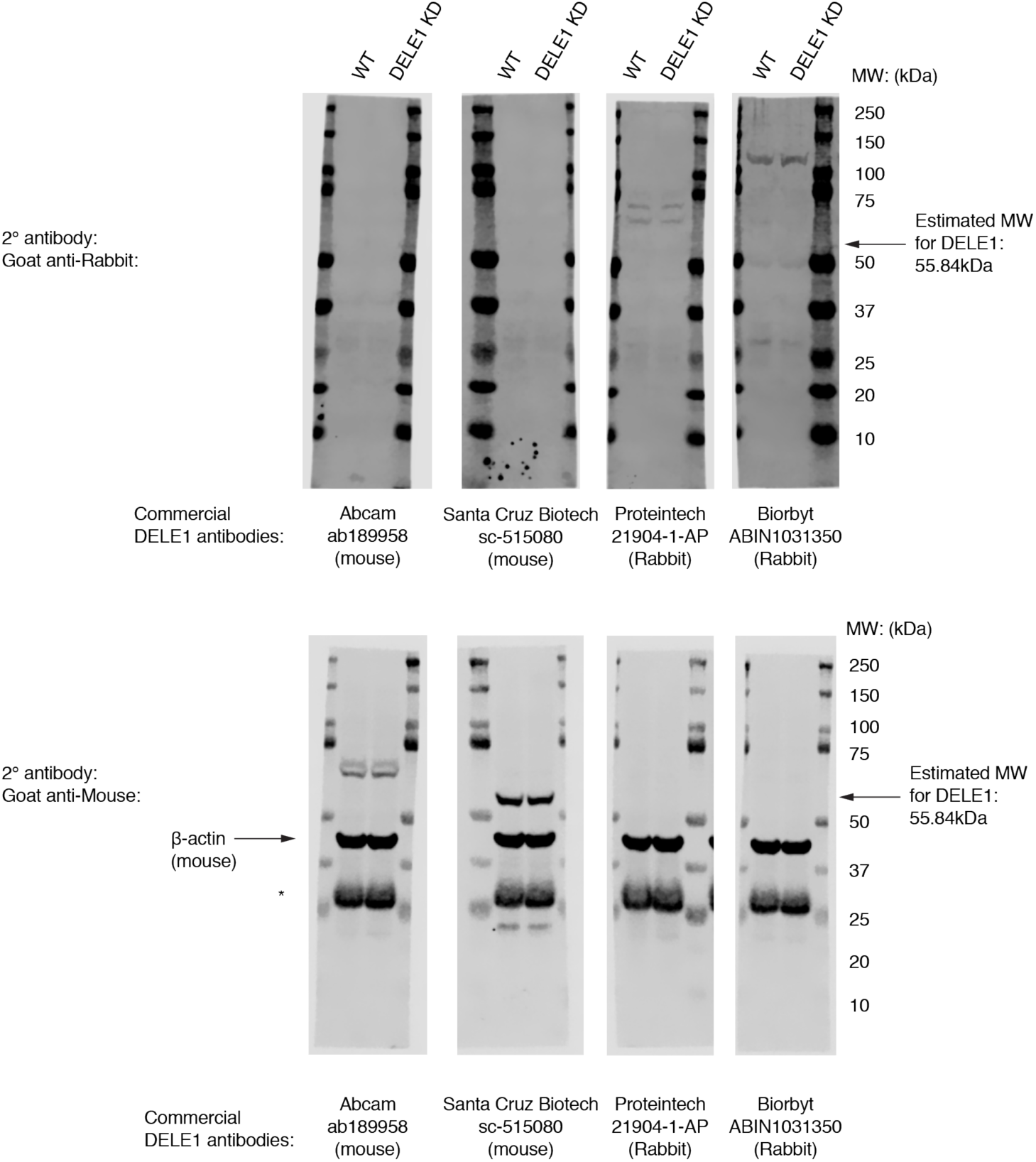
Commercially available antibodies fail to detect DELE1. Lysates from HEK293T cells that were either WT or expressing a sgRNA knocking down DELE1 were probed with the indicated DELE1 antibodies and an antibody against β-actin. *Top*, signal with secondary anti-rabbit antibody. *Bottom*, signal with secondary anti-mouse antibody. None of the bands detected by the DELE1 antibodies decreases in intensity in DELE1 knockdown cells. *Non-specific band.

**Extended Data Figure 5.**
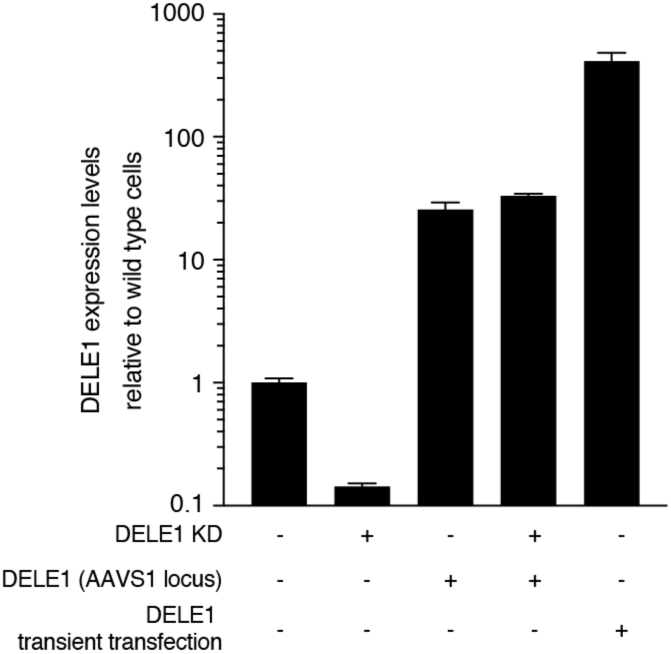
Expression levels of DELE1 in knockdown and overexpression cells. Quantitative RT-PCR quantification of DELE1 mRNA levels in HEK293T cells knocking down (KD) DELE1 by CRISPRi and/or expressing DELE1-mClover stably from the AAVS1 safe-harbor locus or via transient transfection (mean ± s.d., *n* =3 technical replicates).

**Extended Data Figure 6.**
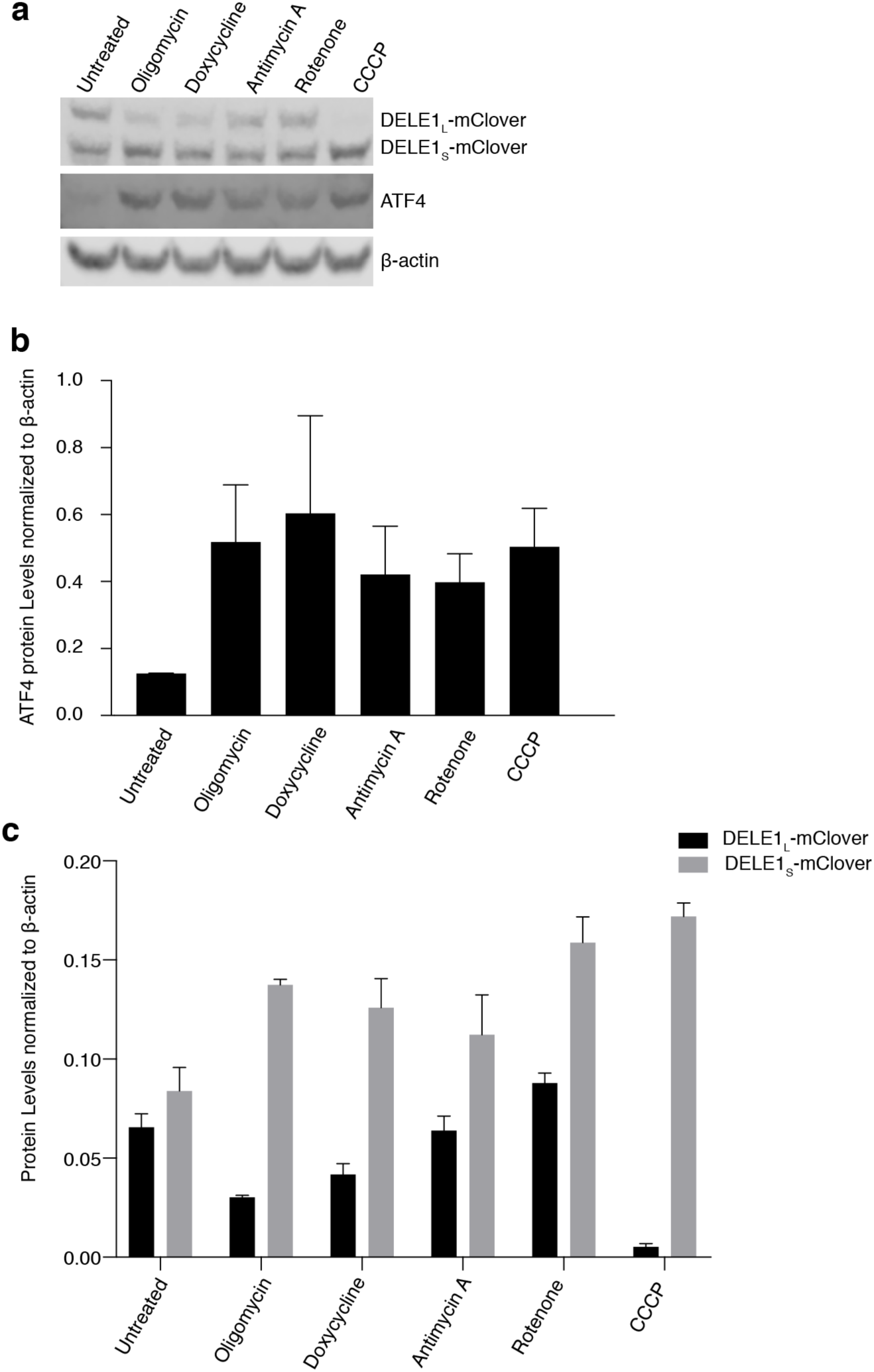
A broad range of mitochondrial toxins stimulates the accumulation of DELE1_S_. Cells stably expressed DELE1-mClover were untreated or treated with a panel of mitochondrial toxins for 16 h (see Methods for details), and subjected to Western blotting with antibodies detecting DELE1-mClover, ATF4 and actin. **(a)** Representative Western blot. **(b)** Quantification of ATF4 levels (mean ± s.d., *n* = 2 blots). **(c)** Quantification of DELE1_L_-mClover and DELE1_S_-mClover levels (mean ± s.d., *n* = 2 blots).

**Extended Data Figure 7.**
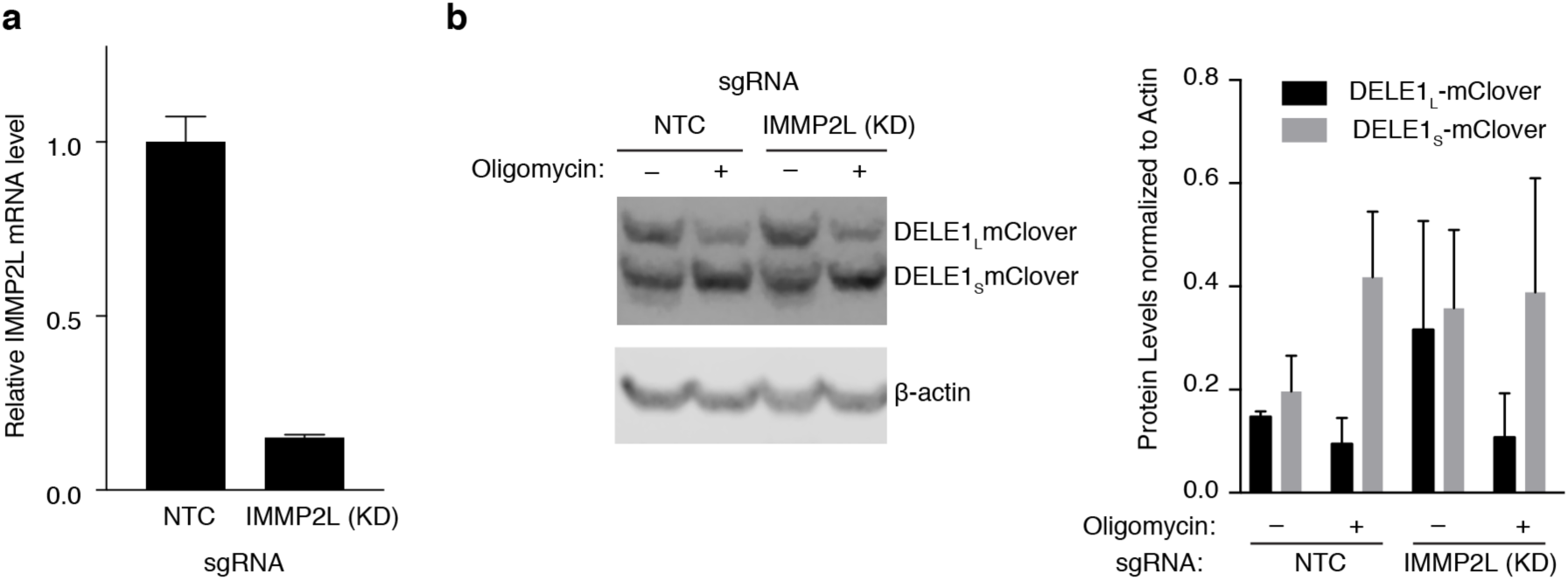
IMMP2L knockdown does not abrogate DELE1 cleavage. **(a)** Quantitative RT-PCR quantification of IMMP2L mRNA levels in HEK293T cells knocking down (KD) IMMP2L by CRISPRi or expressing a non-targeting control (NTC) sgRNA. Mean ± s.d., *n* =3 technical replicates. **(b)** *Left*, Representative Western blot of DELE1-mClover and β-actin in cells expressing non-targeting control sgRNAs (NTC) or sgRNAs for IMMP2L knockdown, which were treated with 1.25 ng/mL of oligomycin for 16 h or left untreated. *Right*, quantification of *n* = 2 blots (mean ± s.d.).

**Extended Data Figure 8.**
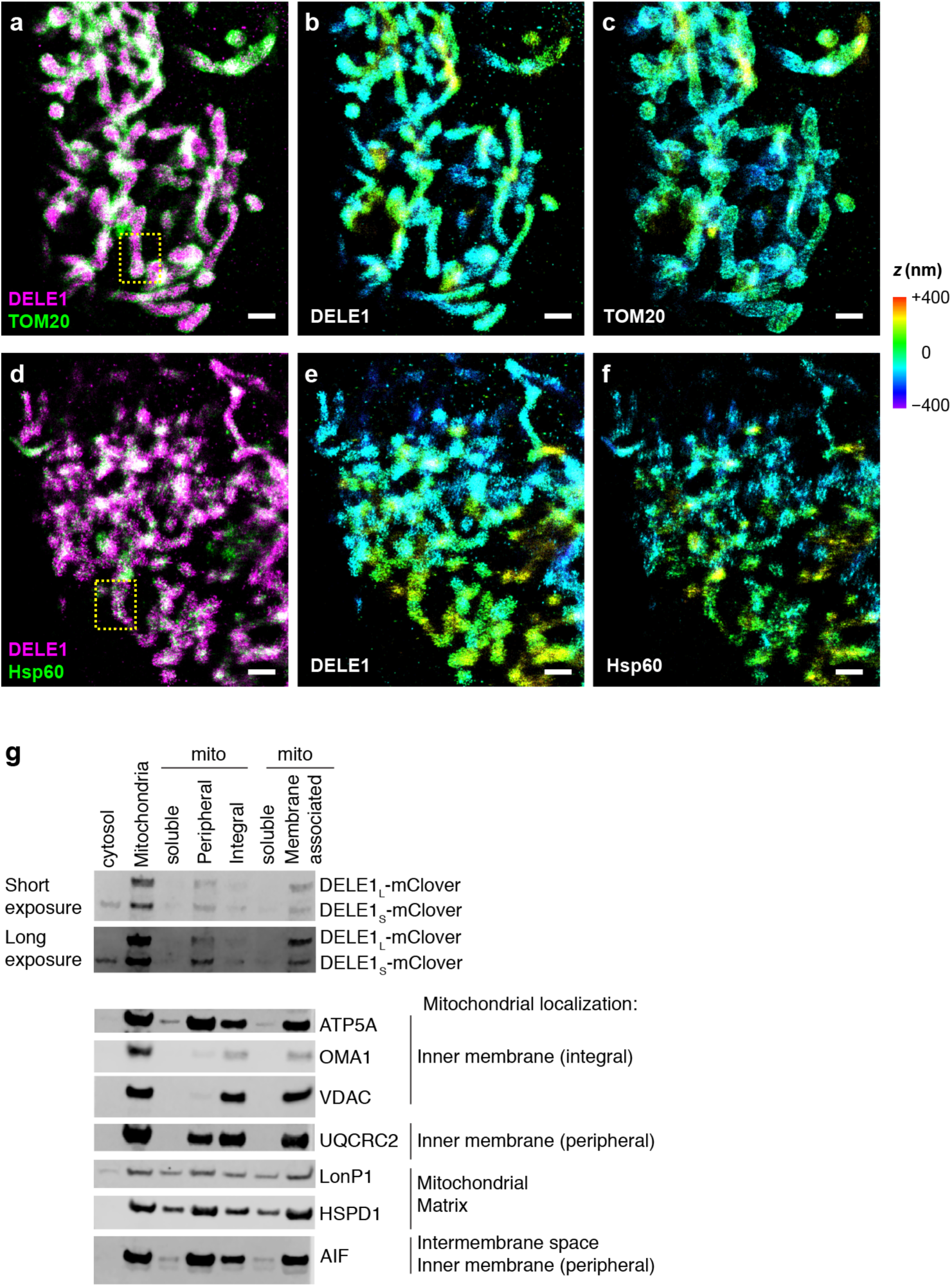
Characterization of DELE1 submitochondrial localization. (**a-f**) Zoom-out views for two-color 3D-STORM super-resolution images of DELE1-mClover vs. TOM20 and Hsp60. Scale bars: 1 µm. (a) Two-color DELE1-mClover (magenta) vs. TOM20 (green). (b,c) The two separated color channels, colored by depth (*Z*). (d) Two-color DELE1-mClover (magenta) vs. Hsp60 (green). (e,f) The two separated color channels, colored by depth (*Z*). The boxed regions in (a,d) correspond to Fig. 3h. (**g**) Biochemical fractionation indicates that DELE1 associates with mitochondrial membranes. Cells stably expressing DELE1-mClover were fractionated into cytosol and mitochondria. The mitochondria were further separated into a soluble and a membrane-associated fraction, or peripheral membrane proteins were extracted using sodium carbonate to differentiate them from integral membrane proteins. Marker proteins with known submitochondrial localizations indicate that the fractionation was not cleanly separating the different types of proteins. However, the soluble mitochondrial fraction contains known matrix and intermembrane space proteins (LonP1, HSPD1, AIF) whereas integral and peripheral membrane proteins of the inner membrane are absent from the soluble fraction. Similarly, DELE1_L_-mClover is not detected in the soluble fraction, suggesting that it is associated with mitochondrial membranes.

**Extended Data Figure 9.**
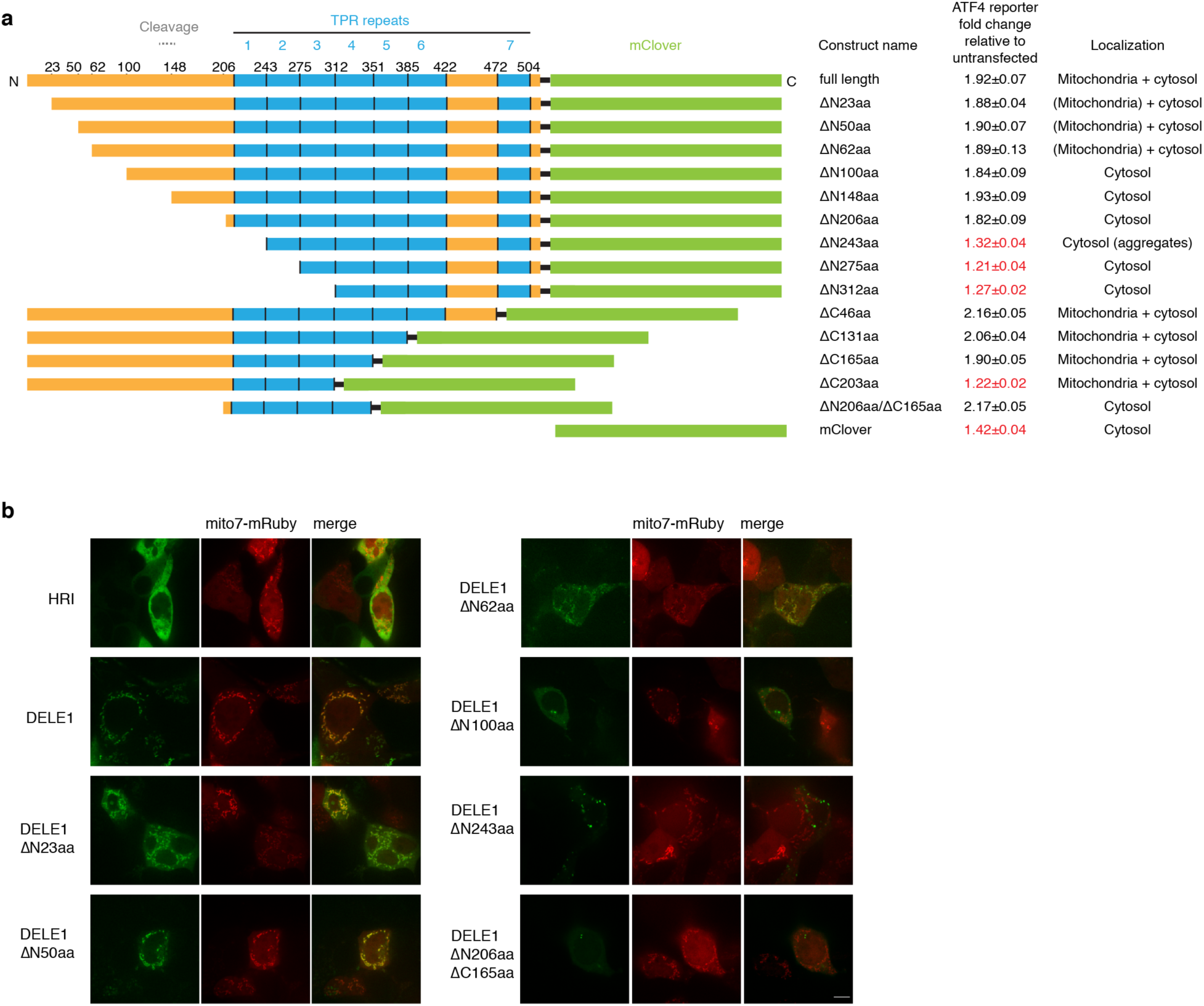
Structure-function analysis of DELE1. **(a)** The indicated DELE1-mClover constructs were transiently overexpressed in reporter cells, and reporter induction was quantified by flow cytometry. Subcellular localization was evaluated by microscopy in cells also expressing mitochondrial-targeted mRuby (b). **(b)** Co-localization of transiently expressed DELE1-mClover with the mitochondrial-targeted mRuby (Mito7-mRuby). Scale bar, 7 μm.

**Extended Data Figure 10.**
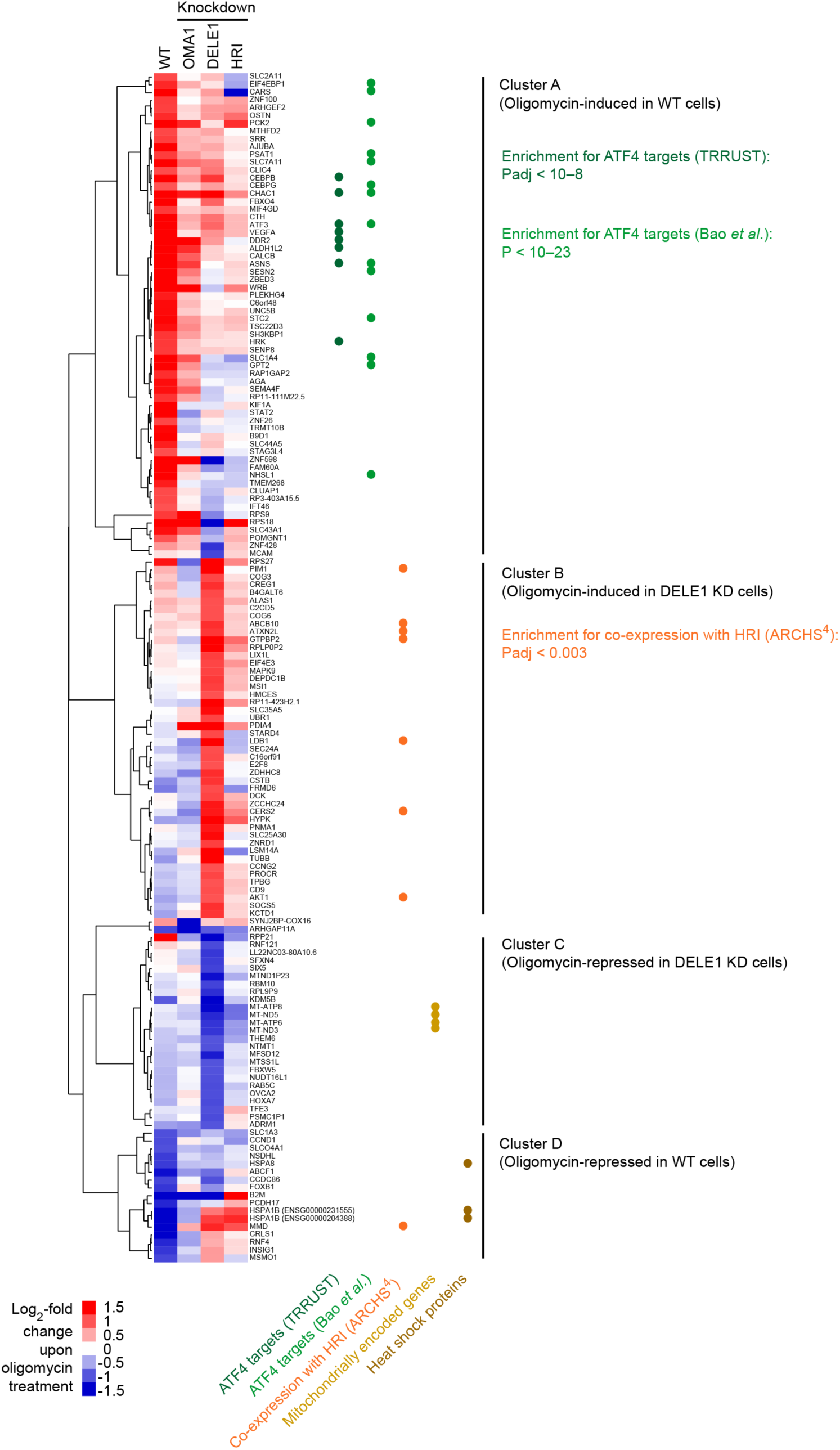
Version of Fig. 5c with genes labeled.

**Extended Data Table 1.**
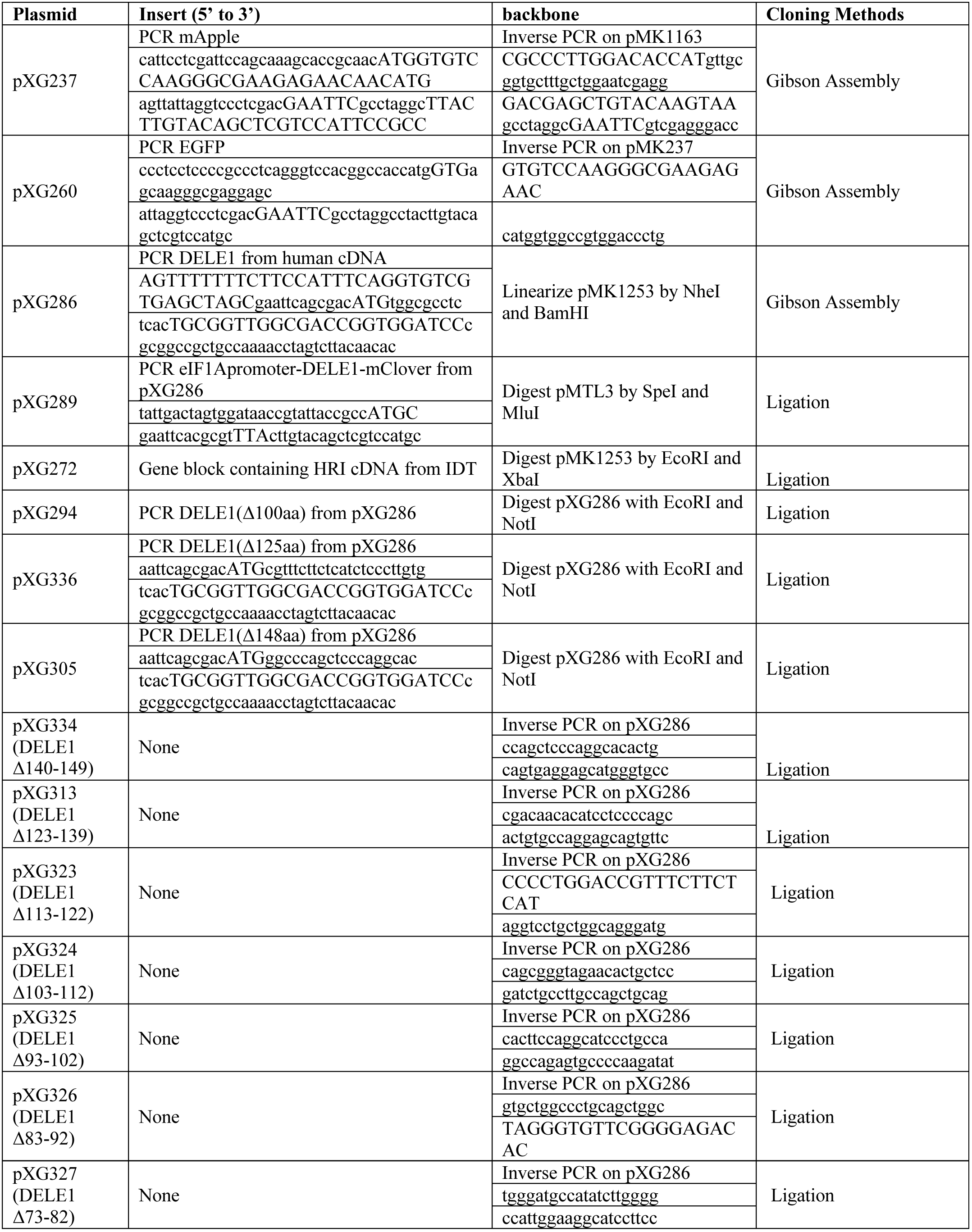
Cloning strategies and oligonucleotide sequences used for plasmids generated.

**Extended Data Table 2.**
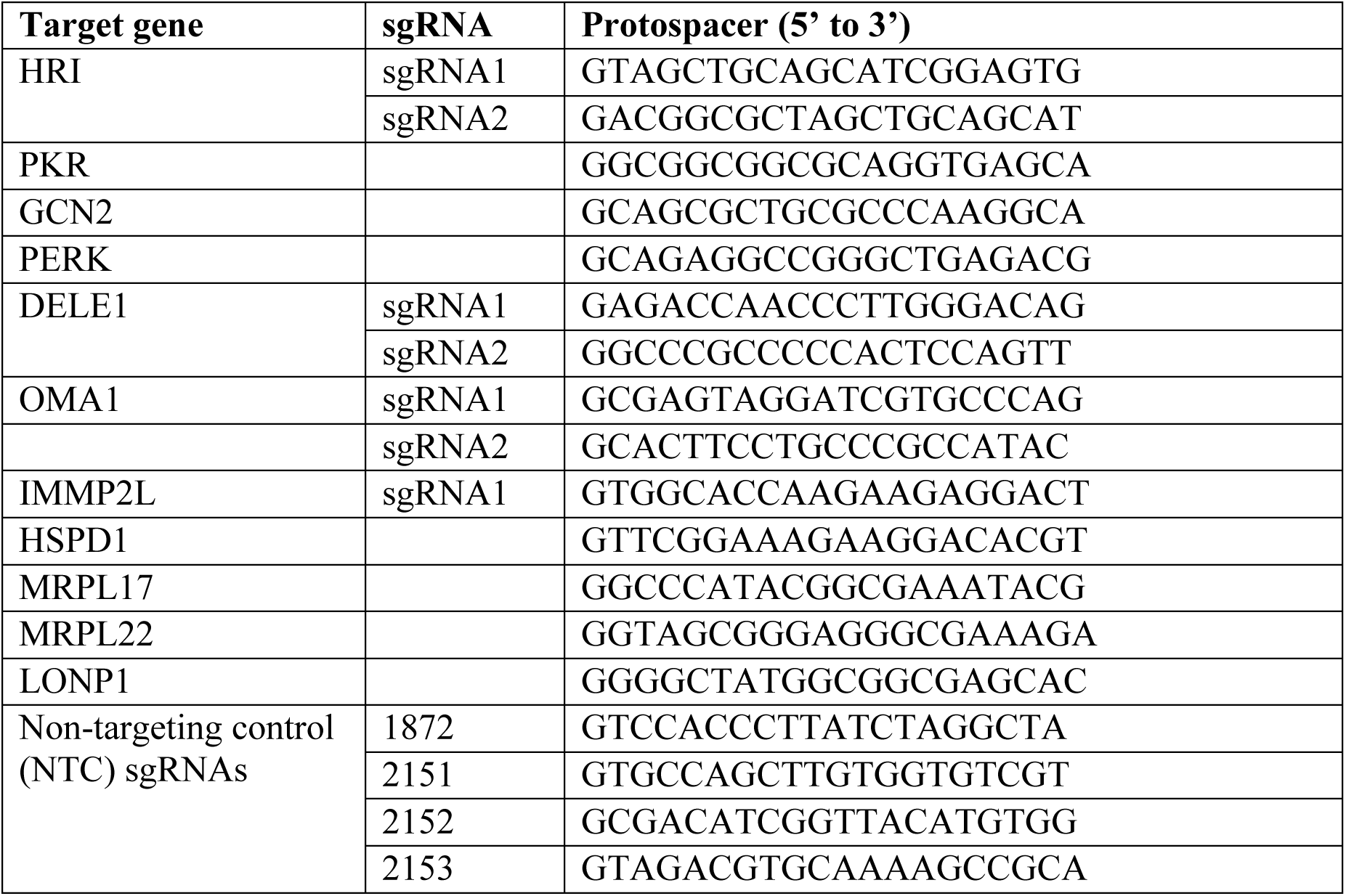
Protospacer sequences of individually cloned sgRNAs.

**Extended Data Table 3.**
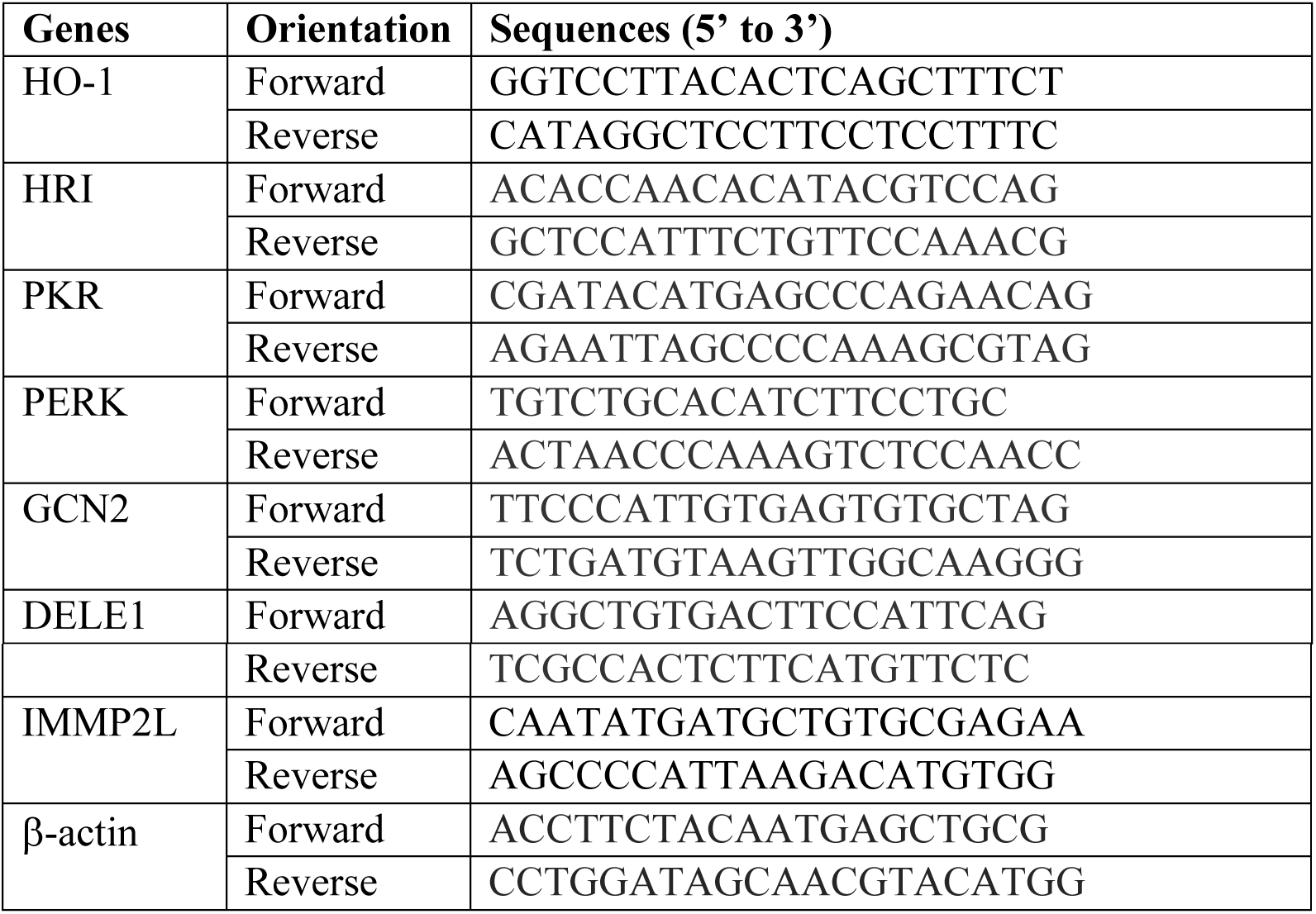
qPCR primer sequences.

## SUPPLEMENTARY TABLE INFORMATION

**Supplemental Tables 1**,**2**,**4-6: Differentially expressed genes based on RNA Sequencing** Differentially expressed genes were determined using DESeq2 (1.20.0)^60^ (see Methods for details). Columns are: A: Gene Identifiers, B: mean normalized counts, averaged over all samples from both conditions, C: log_2_-fold change, D: standard error estimate for the log_2_-fold change estimate, E: Wald statistic, F: P value for this change, G: P value adjusted for multiple testing using the Benjamini-Hochberg procedure which controls false discovery rate.

Supplemental Table 1: Differentially expressed genes upon doxycycline treatment in WT HEK293T cells

Supplemental Table 2: Differentially expressed genes upon oligomycin treatment in WT HEK293T cells

Supplemental Table 4: Differentially expressed genes upon oligomycin treatment in OMA1 KD HEK293T cells

Supplemental Table 5: Differentially expressed genes upon oligomycin treatment in DELE1 KD HEK293T cells

Supplemental Table 6: Differentially expressed genes upon oligomycin treatment in HRI KD HEK293T cells

**Supplemental Table 3: Results from the CRISPRi screen**

Phenotypes for genes targeted in the CRISPRi screens for untreated and oligomycin-treated cells were analyzed using MAGeCK-iNC^26^ (see Methods for details). Columns are: A: Targeted transcription start site, B: targeted gene, C: Knockdown phenotype for screen in oligomycin-treated cells, D: P value for screen in oligomycin-treated cells, E: Gene score = product of phenotype x –log_10_(P value) for screen in oligomycin-treated cells, F: Knockdown phenotype for screen in untreated cells, G: P value for screen in untreated cells, H: Gene score = product of phenotype x –log_10_(P value) for screen in untreated cells.

## References

1 Mattson, M. P., Gleichmann, M. & Cheng, A. Mitochondria in neuroplasticity and neurological disorders. Neuron 60, 748–766, doi:10.1016/j.neuron.2008.10.010 (2008).

2 Sun, N., Youle, R. J. & Finkel, T. The Mitochondrial Basis of Aging. Molecular cell 61, 654–666, doi:10.1016/j.molcel.2016.01.028 (2016).

3 Suomalainen, A. & Battersby, B. J. Mitochondrial diseases: the contribution of organelle stress responses to pathology. Nature reviews. Molecular cell biology 19, 77–92, doi:10.1038/nrm.2017.66 (2018).

4 Nargund, A. M., Pellegrino, M. W., Fiorese, C. J., Baker, B. M. & Haynes, C. M. Mitochondrial import efficiency of ATFS-1 regulates mitochondrial UPR activation. Science 337, 587–590, doi:10.1126/science.1223560 (2012).

5 Fiorese, C. J. et al. The Transcription Factor ATF5 Mediates a Mammalian Mitochondrial UPR. Curr Biol 26, 2037–2043, doi:10.1016/j.cub.2016.06.002 (2016).

6 Munch, C. The different axes of the mammalian mitochondrial unfolded protein response. BMC Biol 16, 81, doi:10.1186/s12915-018-0548-x (2018).

7 Munch, C. & Harper, J. W. Mitochondrial unfolded protein response controls matrix pre-RNA processing and translation. Nature 534, 710–713, doi:10.1038/nature18302 (2016).

8 Zhao, Q. et al. A mitochondrial specific stress response in mammalian cells. The EMBO journal 21, 4411–4419, doi:10.1093/emboj/cdf445 (2002).

9 Papa, L. & Germain, D. SirT3 regulates the mitochondrial unfolded protein response. Mol Cell Biol 34, 699–710, doi:10.1128/MCB.01337-13 (2014).

10 Papa, L. & Germain, D. Estrogen receptor mediates a distinct mitochondrial unfolded protein response. J Cell Sci 124, 1396–1402, doi:10.1242/jcs.078220 (2011).

11 Bao, X. R. et al. Mitochondrial dysfunction remodels one-carbon metabolism in human cells. eLife 5, doi:10.7554/eLife.10575 (2016).

12 Quiros, P. M. et al. Multi-omics analysis identifies ATF4 as a key regulator of the mitochondrial stress response in mammals. J Cell Biol 216, 2027–2045, doi:10.1083/jcb.201702058 (2017).

13 Viader, A. et al. Aberrant Schwann cell lipid metabolism linked to mitochondrial deficits leads to axon degeneration and neuropathy. Neuron 77, 886–898, doi:10.1016/j.neuron.2013.01.012 (2013).

14 Harding, H. P. et al. An integrated stress response regulates amino acid metabolism and resistance to oxidative stress. Molecular cell 11, 619–633 (2003).

15 Wek, R. C., Jiang, H. Y. & Anthony, T. G. Coping with stress: eIF2 kinases and translational control. Biochem Soc Trans 34, 7–11, doi:10.1042/BST20060007 (2006).

16 Taniuchi, S., Miyake, M., Tsugawa, K., Oyadomari, M. & Oyadomari, S. Integrated stress response of vertebrates is regulated by four eIF2alpha kinases. Sci Rep 6, 32886, doi:10.1038/srep32886 (2016).

17 Wek, R. C. Role of eIF2alpha Kinases in Translational Control and Adaptation to Cellular Stress. Cold Spring Harb Perspect Biol 10, doi:10.1101/cshperspect.a032870 (2018).

18 Qi, L. S. et al. Repurposing CRISPR as an RNA-guided platform for sequence-specific control of gene expression. Cell 152, 1173–1183, doi:10.1016/j.cell.2013.02.022 (2013).

19 Gilbert, L. A. et al. Genome-Scale CRISPR-Mediated Control of Gene Repression and Activation. Cell 159, 647–661, doi:10.1016/j.cell.2014.09.029 (2014).

20 Chen, J. J. & London, I. M. Regulation of protein synthesis by heme-regulated eIF-2 alpha kinase. Trends Biochem Sci 20, 105–108 (1995).

21 Kafina, M. D. & Paw, B. H. Intracellular iron and heme trafficking and metabolism in developing erythroblasts. Metallomics 9, 1193–1203, doi:10.1039/c7mt00103g (2017).

22 Lu, L., Han, A. P. & Chen, J. J. Translation initiation control by heme-regulated eukaryotic initiation factor 2alpha kinase in erythroid cells under cytoplasmic stresses. Mol Cell Biol 21, 7971–7980, doi:10.1128/MCB.21.23.7971-7980.2001 (2001).

23 Matts, R. L., Schatz, J. R., Hurst, R. & Kagen, R. Toxic heavy metal ions activate the heme-regulated eukaryotic initiation factor-2 alpha kinase by inhibiting the capacity of hemin-supplemented reticulocyte lysates to reduce disulfide bonds. The Journal of biological chemistry 266, 12695–12702 (1991).

24 Horlbeck, M. A. et al. Compact and highly active next-generation libraries for CRISPR-mediated gene repression and activation. eLife 5, doi:10.7554/eLife.19760 (2016).

25 Kampmann, M., Bassik, M. C. & Weissman, J. S. Integrated platform for genome-wide screening and construction of high-density genetic interaction maps in mammalian cells. Proceedings of the National Academy of Sciences of the United States of America 110, E2317–2326, doi:10.1073/pnas.1307002110 (2013).

26 Tian, R. et al. CRISPR-based platform for multimodal genetic screens in human iPSC-derived neurons. bioRxiv 513309; doi: https://doi.org/10.1101/513309 (2019).

27 Harada, T., Iwai, A. & Miyazaki, T. Identification of DELE, a novel DAP3-binding protein which is crucial for death receptor-mediated apoptosis induction. Apoptosis 15, 1247–1255, doi:10.1007/s10495-010-0519-3 (2010).

28 Baker, M. J. et al. Stress-induced OMA1 activation and autocatalytic turnover regulate OPA1-dependent mitochondrial dynamics. The EMBO journal 33, 578–593, doi:10.1002/embj.201386474 (2014).

29 Ehses, S. et al. Regulation of OPA1 processing and mitochondrial fusion by m-AAA protease isoenzymes and OMA1. J Cell Biol 187, 1023–1036, doi:10.1083/jcb.200906084 (2009).

30 Kim, H., Botelho, S. C., Park, K. & Kim, H. Use of carbonate extraction in analyzing moderately hydrophobic transmembrane proteins in the mitochondrial inner membrane. Protein Sci 24, 2063–2069, doi:10.1002/pro.2817 (2015).

31 Berlanga, J. J., Herrero, S. & de Haro, C. Characterization of the hemin-sensitive eukaryotic initiation factor 2alpha kinase from mouse nonerythroid cells. The Journal of biological chemistry 273, 32340–32346, doi:10.1074/jbc.273.48.32340 (1998).

32 Delaunay, J., Ranu, R. S., Levin, D. H., Ernst, V. & London, I. M. Characterization of a rat liver factor that inhibits initiation of protein synthesis in rabbit reticulocyte lysates. Proceedings of the National Academy of Sciences of the United States of America 74, 2264–2268, doi:10.1073/pnas.74.6.2264 (1977).

33 Walter, P. & Ron, D. The unfolded protein response: from stress pathway to homeostatic regulation. Science 334, 1081–1086, doi:10.1126/science.1209038 (2011).

34 Han, H. et al. TRRUST v2: an expanded reference database of human and mouse transcriptional regulatory interactions. Nucleic acids research 46, D380–D386, doi:10.1093/nar/gkx1013 (2018).

35 Lachmann, A. et al. Massive mining of publicly available RNA-seq data from human and mouse. Nat Commun 9, 1366, doi:10.1038/s41467-018-03751-6 (2018).

36 Abdel-Nour, M. et al. The heme-regulated inhibitor is a cytosolic sensor of protein misfolding that controls innate immune signaling. Science 365, doi:10.1126/science.aaw4144 (2019).

37 Sekine, S. et al. Rhomboid protease PARL mediates the mitochondrial membrane potential loss-induced cleavage of PGAM5. The Journal of biological chemistry 287, 34635–34645, doi:10.1074/jbc.M112.357509 (2012).

38 Uma, S., Hartson, S. D., Chen, J. J. & Matts, R. L. Hsp90 is obligatory for the heme-regulated eIF-2alpha kinase to acquire and maintain an activable conformation. The Journal of biological chemistry 272, 11648–11656, doi:10.1074/jbc.272.17.11648 (1997).

39 Uma, S., Thulasiraman, V. & Matts, R. L. Dual role for Hsc70 in the biogenesis and regulation of the heme-regulated kinase of the alpha subunit of eukaryotic translation initiation factor 2. Mol Cell Biol 19, 5861–5871, doi:10.1128/mcb.19.9.5861 (1999).

40 Michel, S., Canonne, M., Arnould, T. & Renard, P. Inhibition of mitochondrial genome expression triggers the activation of CHOP-10 by a cell signaling dependent on the integrated stress response but not the mitochondrial unfolded protein response. Mitochondrion 21, 58–68, doi:10.1016/j.mito.2015.01.005 (2015).

41 Balsa, E. et al. ER and Nutrient Stress Promote Assembly of Respiratory Chain Supercomplexes through the PERK-eIF2alpha Axis. Molecular cell 74, 877–890 e876, doi:10.1016/j.molcel.2019.03.031 (2019).

42 Park, Y., Reyna-Neyra, A., Philippe, L. & Thoreen, C. C. mTORC1 Balances Cellular Amino Acid Supply with Demand for Protein Synthesis through Post-transcriptional Control of ATF4. Cell Rep 19, 1083–1090, doi:10.1016/j.celrep.2017.04.042 (2017).

43 Khan, N. A. et al. mTORC1 Regulates Mitochondrial Integrated Stress Response and Mitochondrial Myopathy Progression. Cell Metab 26, 419–428 e415, doi:10.1016/j.cmet.2017.07.007 (2017).

44 Chou, A. et al. Inhibition of the integrated stress response reverses cognitive deficits after traumatic brain injury. Proceedings of the National Academy of Sciences of the United States of America 114, E6420–E6426, doi:10.1073/pnas.1707661114 (2017).

45 Das, I. et al. Preventing proteostasis diseases by selective inhibition of a phosphatase regulatory subunit. Science 348, 239–242, doi:10.1126/science.aaa4484 (2015).

46 Acin-Perez, R. et al. Ablation of the stress protease OMA1 protects against heart failure in mice. Sci Transl Med 10, doi:10.1126/scitranslmed.aan4935 (2018).

47 Sidrauski, C. et al. Pharmacological dimerization and activation of the exchange factor eIF2B antagonizes the integrated stress response. eLife 4, e07314, doi:10.7554/eLife.07314 (2015).

48 Chen, J. J. et al. Compromised function of an ESCRT complex promotes endolysosomal escape of tau seeds and propagation of tau aggregation. bioRxiv 637785: doi: https://doi.org/10.1101/637785 (2019).

49 Adamson, B. et al. A Multiplexed Single-Cell CRISPR Screening Platform Enables Systematic Dissection of the Unfolded Protein Response. Cell 167, 1867–1882 e1821, doi:10.1016/j.cell.2016.11.048 (2016).

50 Nagy, T. & Kampmann, M. CRISPulator: A Discrete Simulation Tool For Pooled Genetic Screens. bioRxiv, doi:https://doi.org/10.1101/119131 (2017).

51 Schindelin, J. et al. Fiji: an open-source platform for biological-image analysis. Nat Methods 9, 676–682, doi:10.1038/nmeth.2019 (2012).

52 Rust, M. J., Bates, M. & Zhuang, X. Sub-diffraction-limit imaging by stochastic optical reconstruction microscopy (STORM). Nat. Methods 3, 793–795, doi:10.1038/nmeth929 (2006).

53 Huang, B., Wang, W., Bates, M. & Zhuang, X. Three-dimensional super-resolution imaging by stochastic optical reconstruction microscopy. Science 319, 810–813, doi:10.1126/science.1153529 (2008).

54 Wojcik, M., Hauser, M., Li, W., Moon, S. & Xu, K. Graphene-enabled electron microscopy and correlated super-resolution microscopy of wet cells. Nature Commun. 6, 7384, doi:10.1038/ncomms8384 (2015).

55 Cox, J. & Mann, M. MaxQuant enables high peptide identification rates, individualized p.p.b.-range mass accuracies and proteome-wide protein quantification. Nat Biotechnol 26, 1367–1372, doi:10.1038/nbt.1511 (2008).

56 Liao, M. et al. Impaired dexamethasone-mediated induction of tryptophan 2,3-dioxygenase in heme-deficient rat hepatocytes: translational control by a hepatic eIF2alpha kinase, the heme-regulated inhibitor. J Pharmacol Exp Ther 323, 979–989, doi:10.1124/jpet.107.124602 (2007).

57 Dey, M. et al. PKR and GCN2 kinases and guanine nucleotide exchange factor eukaryotic translation initiation factor 2B (eIF2B) recognize overlapping surfaces on eIF2alpha. Molecular and cellular biology 25, 3063–3075, doi:10.1128/MCB.25.8.3063-3075.2005 (2005).

58 Patro, R., Duggal, G., Love, M. I., Irizarry, R. A. & Kingsford, C. Salmon provides fast and bias-aware quantification of transcript expression. Nat Methods 14, 417–419, doi:10.1038/nmeth.4197 (2017).

59 Soneson, C., Love, M. I. & Robinson, M. D. Differential analyses for RNA-seq: transcript-level estimates improve gene-level inferences. F1000Res 4, 1521, doi:10.12688/f1000research.7563.2 (2015).

60 Love, M. I., Huber, W. & Anders, S. Moderated estimation of fold change and dispersion for RNA-seq data with DESeq2. Genome biology 15, 550, doi:10.1186/s13059-014-0550-8 (2014).

61 Chen, E. Y. et al. Enrichr: interactive and collaborative HTML5 gene list enrichment analysis tool. BMC Bioinformatics 14, 128, doi:10.1186/1471-2105-14-128 (2013).

62 Kuleshov, M. V. et al. Enrichr: a comprehensive gene set enrichment analysis web server 2016 update. Nucleic acids research 44, W90–97, doi:10.1093/nar/gkw377 (2016).

63 Eisen, M. B., Spellman, P. T., Brown, P. O. & Botstein, D. Cluster analysis and display of genome-wide expression patterns. Proceedings of the National Academy of Sciences of the United States of America 95, 14863–14868, doi:10.1073/pnas.95.25.14863 (1998).

64 Saldanha, A. J. Java Treeview--extensible visualization of microarray data. Bioinformatics 20, 3246–3248, doi:10.1093/bioinformatics/bth349 (2004).

